# p53β, an alternatively spliced isoform of p53, induces tumor-suppressive cellular senescence and suppresses tumor growth in wild-type and mutant R273H *TP53* glioblastoma

**DOI:** 10.64898/2026.11.25.769170

**Authors:** Natalia von Muhlinen, Jessica A. Beck, Sebastien Joruiz, Shinji Nakamichi, Chenran Zhang, Kyra Ungerleider, Akriti Trehan, Izumi Horikawa, Curtis C. Harris

**Author notes:** Corresponding author: NVM. The authors declare no competing financial interests.

## Abstract

Glioblastoma (GBM) is the most common and aggressive primary brain tumor, with approximately 30% of cases harboring *TP53* mutations. *TP53* encodes several isoforms, including p53β, which is negatively regulated by the splicing factor SRSF3 and cooperates with p53α to induce tumor-suppressive cellular senescence in normal cells. However, the role of p53β in glioblastoma remains unknown. Here, we showed that increasing wild-type or mutant R273H p53β expression, via SRSF3 depletion or doxycycline-inducible overexpression, induced cellular senescence in GBM cells. Wild-type or mutant R273H p53β also suppressed proliferation and reduced spheroid growth and invasion in *TP53* wild-type and mutant GBM cells, respectively. Mechanistically, mutant R273H p53β induced *CDKN1A*, *BAX*, and *BBC3* mRNA expression, co-immunoprecipitated p53α R273H, and restored p21 promoter activity in a p53α-dose-dependent manner. Transcriptomic analysis revealed distinct gene expression profiles induced by wild-type and mutant R273H p53β that converged on repression of oncogenic pathways that otherwise drive cell cycle and proliferation in GBM. In orthotopic xenografts, wild-type and mutant R273H p53β reduced tumor growth and prolonged event-free survival. These findings identify p53β as a potential therapeutic target in WT and R273-mutant *TP53* GBM and show that a mutant p53 isoform can redirect an oncogenic *TP53* hotspot mutant toward tumor suppression.

## Introduction

Glioblastoma (GBM), an *IDH*-wild-type, CNS WHO grade 4 diffuse astrocytic glioma, is the most common and aggressive primary malignant brain cancer in adults [1, 2]. *TP53,* a central regulator of stress responses and cell fate [3], is mutated in approximately 30% of GBMs [4]. In *TP53* wild-type (WT) tumors, p53 function can also be impaired by suppressed *TP53* expression [5], dysregulated p53 factors, and impaired p53 target gene induction [4, 6–8]. Restoring p53-mediated tumor-suppressive functions, including cellular senescence [9], may therefore represent a therapeutic strategy for GBM.

*TP53* encodes at least 12 naturally occurring isoforms, including full-length p53 (p53α) through alternative RNA splicing, promoter usage, and translation initiation [10, 11]. These isoforms modulate canonical p53α activity and may contribute to p53 pathway dysregulation in cancer [3]. We previously reported that the C-terminally truncated p53 isoform, p53β [10], cooperates with p53α to promote senescence in normal human cells [12–14], and that its negative regulation by serine/arginine-rich splicing factor 3 (SRSF3) via alternative splicing promotes senescence [14, 15]. SRSF3 is frequently upregulated in cancers, including GBM [16, 17], but whether it regulates p53β-mediated senescence in cancer remains unknown.

Whether p53β retains tumor-suppressive activity when it harbors somatic *TP53* mutations remains unexplored. R273H, among the most frequent *TP53* hotspot mutations, disrupts sequence-specific DNA binding and canonical transcriptional activity while largely preserving the overall structure of the DNA-binding domain [18]. Beyond loss of tumor-suppressive activity, R273H confers gain-of-function properties that promote cancer progression [19, 20]. Although the DNA-binding domain, including sites of hotspot mutations, is conserved in most p53 isoforms [10, 11], including p53β, the functional consequences of these mutations have been studied predominantly in p53α.

Here, we used in vitro and orthotopic GBM mouse models to investigate whether p53β expression, through SRSF3 depletion or lentiviral overexpression, retains tumor-suppressive activity in WT and mutant R273H *TP53* backgrounds. Overall, our findings identify an isoform-based mechanism through which an oncogenic *TP53* hotspot mutation retains tumor-suppressive activity in the p53β isoform.

## Results

### SRSF3 depletion increases p53β isoform expression and induces senescence and apoptosis in WT and R273H mutant *TP53* GBM cells

We re-analyzed TCGA glioma datasets using the 2021 WHO classification to examine *SRSF3* mRNA expression in gliomas [21]. *SRSF3* expression was significantly elevated in *IDH-*wild-type GBM compared with lower-grade glioma (LGG) and normal brain tissue **(Supplementary Fig. 1A)**, consistent with previous reports [16, 17]. *SRSF3* expression did not differ significantly between *TP53* wild-type and mutant gliomas **(Supplementary Fig. 1B,C)**, suggesting that its upregulation is independent of *TP53* status.

To determine whether SRSF3 regulates p53β expression in GBM, we depleted SRSF3 using two independent siRNAs [15] in U87-MG (*TP53* WT) and SF268 (*TP53* R273H). SRSF3 knockdown increased p53β mRNA and protein expression **(Supplementary Fig. 1D,E)**, increased senescence-associated-β-galactosidase (SA-β-gal) activity **(Supplementary Fig. 1F,G)**, decreased Ki67 staining, and increased cleaved caspase 3 (CC3) **(Supplementary Fig. 1H,I)** in both GBM cell lines. SRSF3 inhibition also increased expression of the canonical p53 target genes *CDKN1A* (p21), *BAX* and *BBC3* (*PUMA*) in WT and mutant R273H *TP53* GBM cells **(Supplementary Fig. 1J,K)**.

To assess whether cellular senescence following SRSF3 depletion was dependent on expression of p53 or its isoforms, we depleted SRSF3 in LNZ308N (*TP53* null) cells. Unlike U87-MG and SF268, which express endogenous p53, LNZ308N cells did not show increased SA-β-gal or induction of the assessed p53 target genes following SRSF3 depletion **(Supplementary Fig. 1L-N)**. Together, these results show that SRSF3 depletion increases p53β expression and induces senescence and apoptosis-associated responses in both *TP53* WT and R273H GBM cells in a *TP53*-dependent manner.

### Mutant p53β R273H expression suppresses proliferation and induces cellular senescence in *TP53* R273H GBM cells

To examine the functional effects of p53β expression, we generated GBM cell lines stably expressing doxycycline-inducible (Tet-On) WT p53β (U87-MG and A172; *TP53* WT), mutant p53β R273H (SNB19, SF268, *TP53* R273H), or carrying an empty-vector (EV) control **(Fig. 1A)**. Endogenous p53β was undetectable at the mRNA **(Fig. 1B)** or protein **(Fig. 1C)** levels, consistent with the high SRSF3 expression observed in GBM cells **(Supplementary Fig. 1)**, whereas doxycycline (DOX, 0.1 µg/ml) induced robust overexpression of WT or mutant p53β R273H **(Fig. 1B,C)**, in the corresponding cells. Consistent with our findings in normal fibroblasts [15], DOX-induced WT p53β significantly decreased proliferation in U87-MG and A172 cells **(Fig. 1D,E)**. Similarly, expression of mutant p53β R273H, significantly reduced proliferation in SF268 and SNB19 GBM cells, despite harboring the oncogenic R273H mutation **(Fig. 1D,E)**. In contrast, DOX treatment of EV-transduced cells did not affect proliferation **(Fig. 1D).** In addition, expression of WT or mutant R273H p53β, but not an EV control, also significantly increased senescence-associated β-galactosidase activity and p21 immunostaining **(Fig. 1F,G)**.

**Figure 1:**
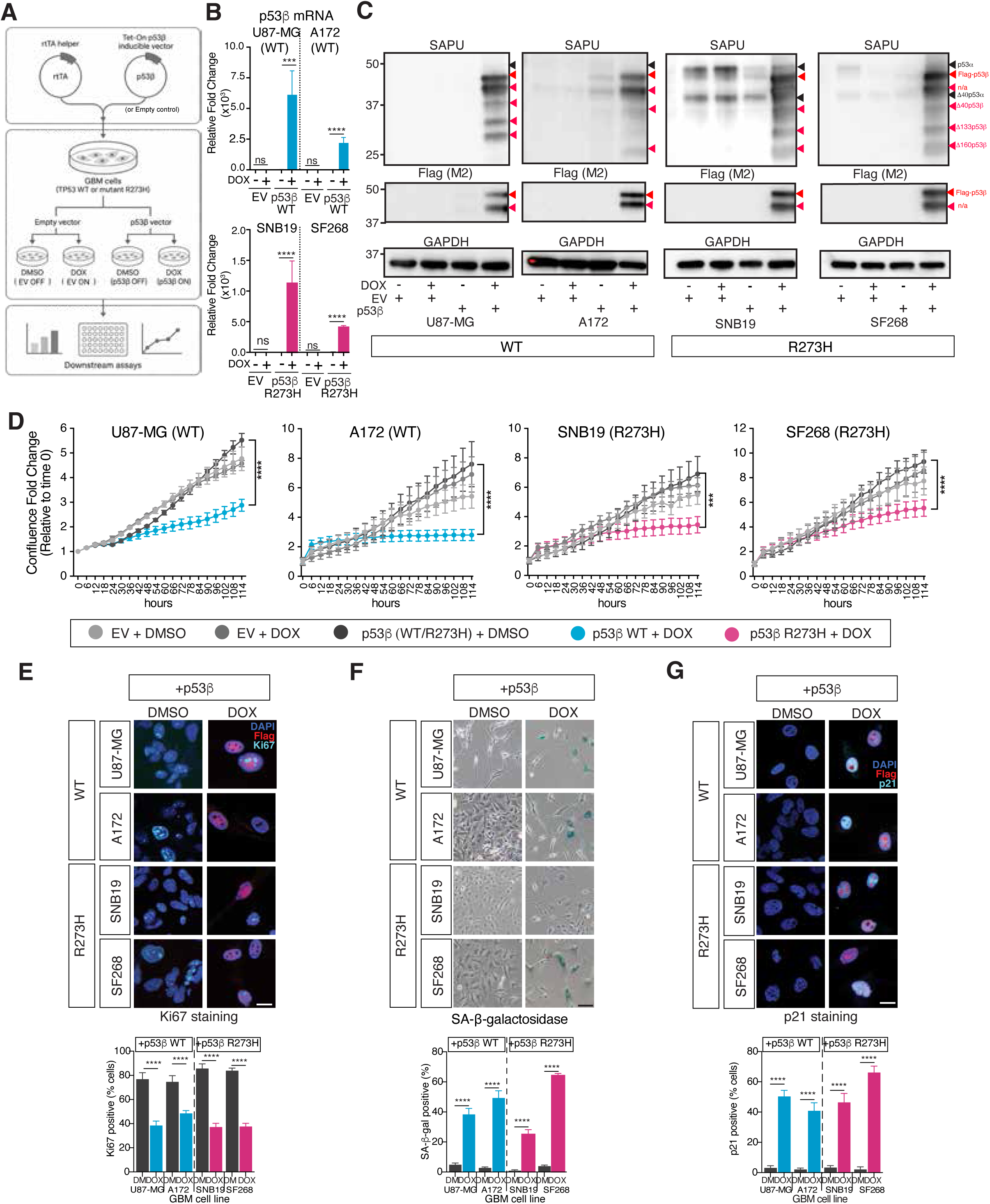
Doxycycline-inducible expression of WT or mutant R273H p53β suppresses proliferation and induces senescence in GBM cells harboring wild-type or mutant R273H *TP53*. **(A)** Schematic of the experimental workflow for doxycycline-inducible expression of WT or mutant R273H *TP53* in GBM cell lines, with empty vector (EV) as a control. **(B)** RT-qPCR analysis of relative p53β mRNA expression normalized to GAPDH. **(C)** Western blot analysis of p53 isoforms (SAPU, a pantromic antibody that detects most p53 isoforms [10]) and Flag-tagged p53β (FLAG M2). GAPDH served as a loading control. Cropped blots are shown; uncropped blots are provided in Supplementary Figure 6. Black arrows: endogenous protein. Red arrows: overexpressed protein. n/a: unidentified. **(D)** Cell confluence (fold change relative to time 0) monitored over 5 days. **(E-G)** Representative images and quantification of the percentage of cells positive for **(E)** Ki67, **(F)** SA-β-gal activity, and (G) p21 in the indicated GBM cell lines expressing inducible WT or mutant R273H p53β. Cells were treated with DMSO or doxycycline (DOX) for 72-96 h (B-G). Scale bars: 100μm (F); 20μm (E,G). Data are presented as mean ± SD from three independent experiments (*n* = 3). Statistical significance was assessed by two-way ANOVA followed by Sidak’s (B, E-G) or Tukey’s (D) multiple comparisons test. *p < 0.05; **p<0.01; ***p<0.001; ****p<0.0001. ns, not significant.

Together, these findings demonstrate that mutant R273H p53β retain the ability to suppress proliferation and induce cellular senescence in GBM cells expressing endogenous p53 isoforms, despite harboring a canonical oncogenic *TP53* mutation.

### Mutant p53β R273H retains transcriptional activity through cooperation with mutant p53α

We next investigated whether mutant p53β R273H retains the ability to regulate canonical p53 target genes in GBM cells expressing the corresponding endogenous p53α isoform. DOX-induced expression of WT or mutant R273H p53β, but not an EV control, increased *CDKN1A* (p21), *BAX*, and *BBC3* (*PUMA*) mRNA expression upon doxycycline incubation in the *TP53* WT (U87-MG and A172) and *TP53* R273H (SF268, SNB19) GBM cell lines **(Fig. 2A,B)**.

**Figure 2:**
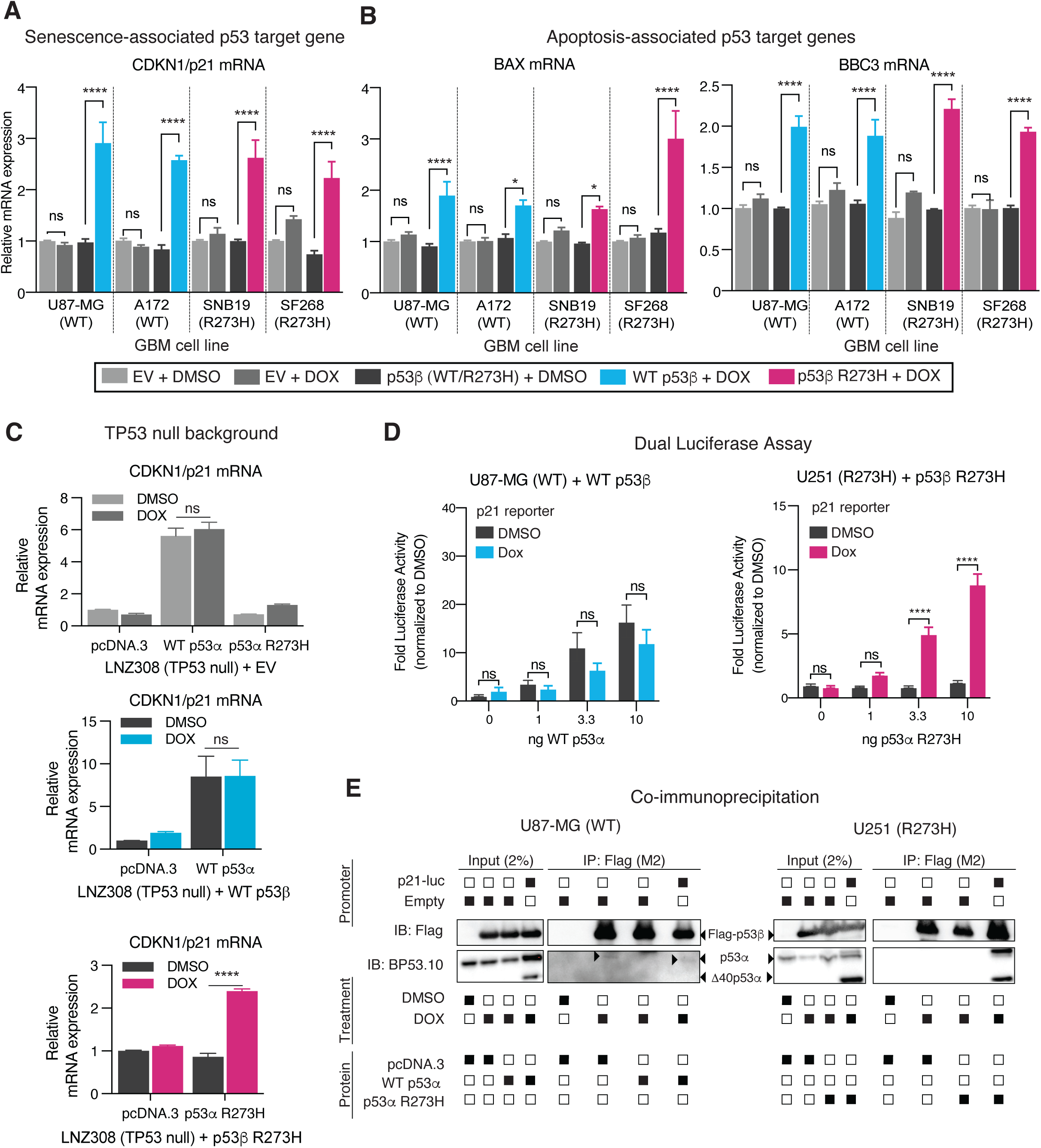
Mutant R273H p53β induces p53 target gene expression and cooperates with mutant p53α R273H to activate the p21 promoter. (A,B) Relative mRNA expression of (A) *CDKN1A* (p21), a senescence-associated p53 target gene, and (B) *BAX* and *BBC3* (*PUMA*), apoptosis-associated p53 target genes, measured by qRT-PCR in the indicated WT or mutant R273H *TP53* GBM cell lines expressing doxycycline-inducible empty vector (EV) or the corresponding p53β isoform following doxycycline (DOX) or DMSO treatment. **(C)** Relative expression of *CDKN1A* (p21) mRNA in *TP53-*null LNZ308N GBM cells stably expressing a doxycycline-inducible EV, WT or mutant R273H p53β, transfected with the indicated plasmids. **(D)** Dual-luciferase reporter assay measuring p21 promoter activity in U87-MG and U251 GBM cells stably expressing doxycycline-inducible WT or mutant R273H p53β, respectively, and transfected with increasing amounts (0-10 ng) of the corresponding p53α expression plasmid. Luciferase activity is expressed as fold change relative to the corresponding DMSO-treated control. **(E)** Co-immunoprecipitation of WT or mutant R273H Flag-tagged p53β in U87-MG and U251 GBM cells, respectively, transfected with the indicated p53α expression or control plasmids, in the presence or absence of the p21 promoter reporter plasmid. Input (2%) and FLAG immunoprecipitates were analyzed by Western blot using the indicated antibodies. Filled and open squares indicate presence or absence, respectively. Cropped blots are shown; uncropped blots are provided in Supplementary Figure 6. Cells were treated with DMSO or DOX (0.1μg/ml) for 96 h (A-C), or overnight (D, E). Relative mRNA expression was normalized to GAPDH **(A-C)**. Data are presented as mean ± SD from three independent experiments (*n*=3). Statistical significance was assessed by two-way ANOVA followed by Sidak’s multiple comparisons test. *p < 0.05; **p<0.01; ***p<0.001; ****p<0.0001. ns, not significant.

Given that these GBM cells express endogenous p53α, we next investigated whether p53β-mediated p53 target induction requires p53α. We expressed WT or mutant R273H p53β, alone or in combination with the corresponding p53α isoform in LNZ308N (*TP53* null) cells. Neither mutant p53β R273H nor mutant p53α R273H alone induced *CDKN1A* expression, whereas co-expression of both mutant isoforms significantly increased *CDKN1A* mRNA levels **(Fig. 2C; Supplementary Fig. 2A-B)**. Thus, mutant p53β R273H requires co-expression of mutant p53α R273H to induce *CDKN1A* in R273H cells.

We assessed p21 promoter activity by dual-luciferase reporter assay in U87-MG (*TP53* WT) and U251 (*TP53* R273H, see Methods) cells, expressing DOX-inducible WT or mutant R273H p53β, respectively. As previously reported [22], WT p53α robustly activated the p21 promoter in a dose-dependent manner, whereas mutant p53α R273H did not **(Fig. 2D)**. Neither WT nor mutant p53β R273H alone activated the p21 promoter. Co-expression of both mutant isoforms significantly increased p21 promoter activity in a p53α-dose-dependent manner **(Fig. 2D)**. Flag-tagged mutant p53β R273H co-immunoprecipitated mutant p53α R273H only in the presence of the p21 promoter-reporter construct **(Fig. 2E)**, consistent with previous findings for WT p53β [22].

Together, these findings demonstrate that, while neither mutant p53β R273H nor mutant p53α R273H alone induces *CDKN1A*, they cooperate to promote *CDKN1A* expression and restore p21 promoter activity. Their association in the presence of a p53-responsive promoter construct supports a mechanism of functional cooperation between the two mutant isoforms, despite both harboring the oncogenic R273H mutation.

### WT and mutant R273H p53β induce distinct gene expression profiles but converge to suppress cell cycle, cell growth, and proliferation pathways

To characterize p53β-induced transcriptional changes, we performed bulk RNA sequencing (RNA-seq) in U87-MG, A172, SF268, and SNB19 cells expressing inducible WT or mutant R273H p53β. Empty-vector (EV) controls were included to assess doxycycline-associated off-target effects **(Supplementary Fig. 3A)**. WT or mutant R273H p53β induction yielded substantially more differentially expressed genes (DEGs; 287-1610 per cell line) than EV controls (7-102 per cell line) **(Fig. 3A-B; Supplementary Fig. 3B-E)**, supporting p53β-dependent transcriptional regulation.

**Figure 3:**
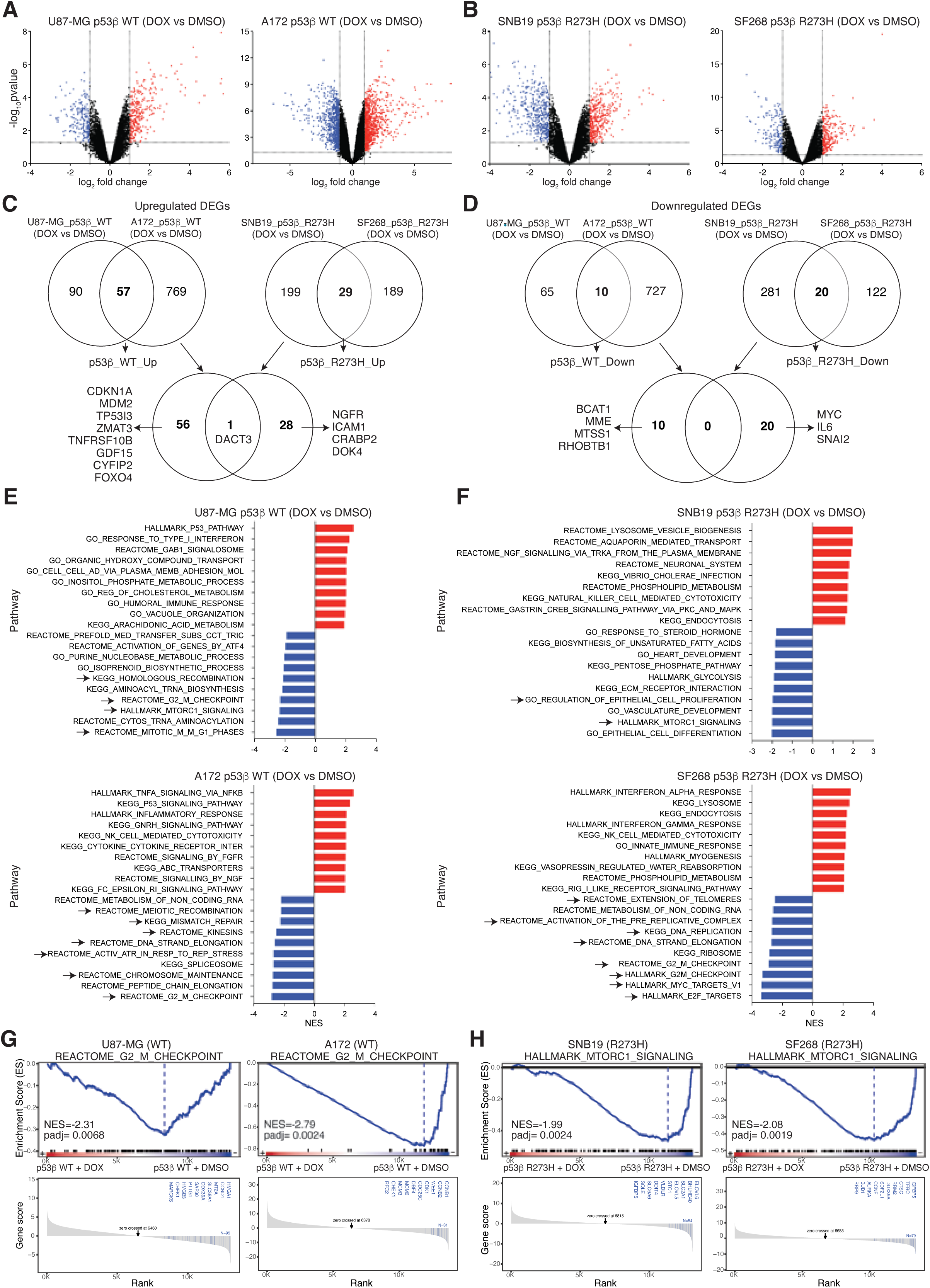
Distinct gene expression profiles in GBM cells stably expressing WT or mutant R273H p53β. RNA sequencing was performed on GBM cell lines stably expressing the corresponding doxycycline-inducible p53β isoform, following treatment with DMSO or DOX (0.1μg/ml) for 48 h, using three independent biological replicates per condition (n=3). **(A,B)** Volcano plots showing log_2_ fold change versus - log_10_(p value) in the indicated WT **(A)** or mutant R273H **(B)** *TP53* GBM cell lines (DOX vs DMSO contrast). Red and blue dots represent significantly up- and downregulated genes, respectively. Dotted lines indicate differential expression thresholds (log_2_FC> | 1 | and p < 0.05). **(C,D)** Venn diagrams representing the upregulated **(C)** and downregulated **(D)** differentially expressed genes (DEGs) within and between GBM cell lines expressing WT or mutant R273H p53β. Selected DEGs are indicated. Labels: p53β_WT_Up and p53β_WT_Down: DEGs commonly up- or downregulated in U87-MG and A172 cells expressing WT p53β; p53β_R273H_Up and p53β_R273H_Down: DEGs commonly up- or downregulated in SNB19 and SF268 cells expressing mutant R273H p53β. **(E,F)** Normalized enrichment scores (NES) of up to 10 top-ranked positively (red) and negatively (blue) enriched gene sets identified by gene set enrichment analysis (GSEA; adjusted p <0.05) in GBM cell lines expressing WT **(E)** or mutant R273H **(F)** p53β. Arrows indicate gene sets related to cell cycle progression, DNA replication/repair, and growth signaling. **(G,H)** GSEA enrichment plots (upper panels) and ranked gene scores (lower panels) for Reactome G2/M checkpoints **(G)** and Hallmark mTORC1 signaling **(H)**, shared by WT and mutant R273H *TP53* GBM cell lines, respectively. NES and adjusted p values are indicated. DEGs and enriched gene sets identified in the corresponding EV DOX vs DMSO contrasts were excluded from all analyses shown.

To identify WT- and R273H-specific transcriptional responses, we compared DEGs shared between the two *TP53* WT (U87-MG and A172) or mutant (SF268 and SNB19) cell lines **(Fig. 3C-D; Supplementary Fig. 3F-G)**. We identified 57 upregulated and 10 downregulated DEGs shared by WT p53β-expressing cells, compared with 29 upregulated and 20 downregulated DEGs shared by mutant p53β R273H GBM cell lines **(Fig. 3C,D)**. Only one gene, *DACT3*, a negative regulator of the Wnt/β-catenin signaling pathway [23, 24], was commonly upregulated by WT and mutant R273H p53β. WT p53β induced predominantly canonical p53 targets, including *CDKN1A*, *MDM2,* and *TP53*I3, while repressing genes including *BCAT1* and *MME.* In contrast, mutant p53β R273H upregulated genes including *NGFR* and *ICAM1*, while repressing oncogenic and mesenchymal regulators such as *MYC*, IL6, and *SNAI2* **(Fig. 3C,D, Supplementary Fig. 3F,G)**. Although *CDKN1A*, *BAX,* and *BBC3* (*PUMA*) were induced by mutant p53β R273H in qRT-PCR **(Fig. 2A-B)**, these genes did not reach significance in the RNA-seq, potentially reflecting the greater sensitivity of qRT-PCR or harvest time (48 h versus 96 h post-DOX).

Gene set enrichment analysis (GSEA) further revealed distinct transcriptional responses to WT and mutant R273H p53β, with substantial heterogeneity among cell lines sharing the same *TP53* status **(Fig. 3E,F)**. Pathway enrichment was therefore evaluated separately for each cell line, excluding pathways significantly enriched in the corresponding empty vector (EV) DOX versus DMSO controls. WT p53β enriched canonical p53 signaling and stress-response pathways while suppressing cell-cycle progression, chromosome maintenance, and growth signaling **(Fig. 3E,G; Supplementary Fig. 3H,I)**. In contrast, mutant p53β R273H suppressed oncogenic mTORC1 and MYC targets (V1) pathways in both SF268 and SNB19, which were the only pathways shared by both cell lines, while modulating pathways associated with interferon/innate immune signaling, endocytosis, and phospholipid metabolism **(Fig. 3F,H; Supplementary Fig. 3J,K)**. L2P pathway analysis further supported these findings and identified enrichment of the HALLMARK p53 pathway in SF268 cells, consistent with partial activation of canonical p53-associated transcription **(Supplementary Fig. 3L,M)**.

Despite differences in individual enriched pathways, the transcriptional responses were consistent with reduced cell-cycle progression and growth signaling, consistent with the shared anti-proliferative phenotypes observed across the four GBM cell lines.

### WT and mutant R273H p53β suppress spheroid growth and invasion in GBM cells

To assess p53β activity in three-dimensional cultures, we generated spheroids from the same GBM cells and U251 (*TP53* R273H), an additional cell line used for subsequent studies (see methods), expressing WT or mutant R273H p53β, or an EV control. WT p53β significantly suppressed spheroid growth in both U87-MG and A172, as did mutant p53β R273H in SF268, SNB19, and U251 **(Fig. 4A,B)**. In contrast, DOX did not alter spheroid growth in EV controls.

**Figure 4:**
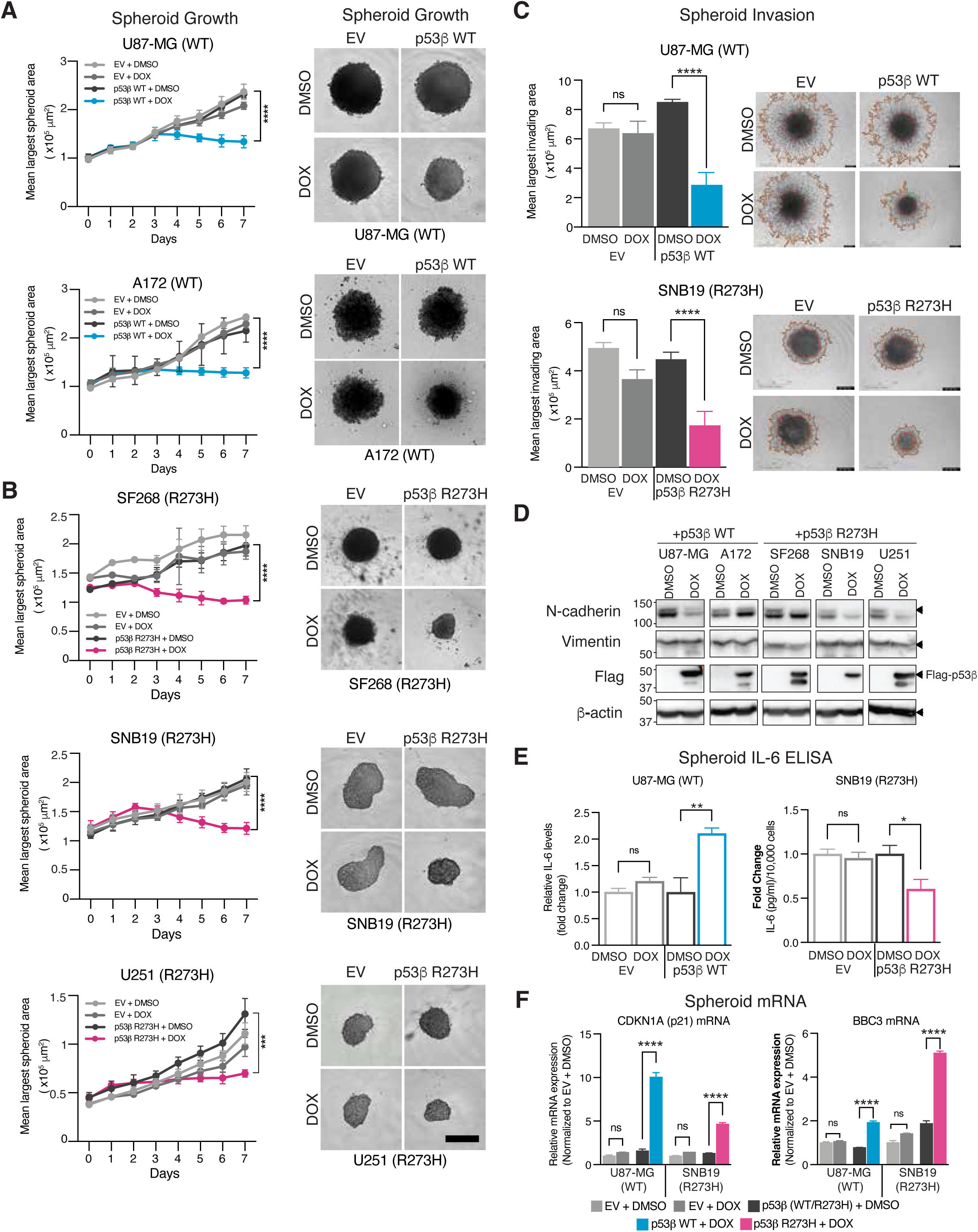
WT or mutant R273H p53β expression suppresses GBM spheroid growth and invasion. (A-C) GBM cell lines expressing a doxycycline-inducible empty vector (EV) or the corresponding p53β vector were plated into ultra-low-attachment 96-well plates, treated with DOX (0.1μg/ml) or DMSO after 24 h incubation, and monitored using an Incucyte S3. **(A,B)** Left: Quantification of the mean largest spheroid area over 7 days in WT **(A)** or mutant R273H **(B)** *TP53* GBM cell lines. DOX was added to the medium on day 2. Right: representative spheroid images at day 7 post-seeding. Scale bar: 400 µm. **(C)** Spheroids were embedded in Matrigel to assess invasion. Left: Quantification of the largest invading area 5 days after Matrigel addition. Right: Representative images of spheroid invasion. Scale bar: 1mm. **(D)** Western blot analysis of the epithelial-to-mesenchymal transition (EMT) markers N-cadherin and vimentin, Flag-tagged p53β, and β-actin as a loading control in conventional 2D GBM cell cultures treated with DOX or DMSO for 96 h. Cropped blots are shown; uncropped blots are provided in Supplementary Figure 6. **(E)** IL-6 quantification by ELISA in spheroid supernatants collected at day 5 post-seeding, normalized to 10,000 cells and expressed as fold change. **(F)** Relative mRNA expression of *CDKN1A* (p21) and *BBC3* (*PUMA*) measured by qRT-PCR in spheroids collected at day 7 post-seeding, normalized to GAPDH and expressed relative to EV + DMSO controls. Data are presented as mean ± SD from three independent experiments (n=3). Statistical significance was assessed by two-way ANOVA **(A,B)** or one-way ANOVA **(C,E,F)** followed by Tukey’s multiple comparisons test. *p < 0.05; **p<0.01; ***p<0.001; ****p<0.0001. ns, not significant.

WT and mutant R273H p53β also significantly reduced spheroid invasion in U87-MG and SNB19 cells, respectively, whereas DOX had no effect in the corresponding EV controls **(Fig. 4C)**. Consistently, p53β expression reduced N-cadherin expression across all five cell lines, while vimentin expression showed no consistent changes **(Fig. 4D)**. These findings are consistent with suppression of HALLMARK_EMT pathway following WT p53β expression in U87-MG cells **(Supplementary Fig. 3L)** and with the downregulation of *SNAI2*, an EMT-associated transcription factor, in mutant p53β R273H-expressing cells **(Fig. 3D)**.

We next investigated whether p53β expression affects interleukin-6 (IL-6) secretion, a cytokine associated with senescence-associated secretory phenotype (SASP) and GBM-associated inflammation. WT p53β expression following DOX treatment significantly increased IL-6 secretion by U87-MG spheroids, whereas mutant p53β R273H significantly decreased IL-6 levels in SNB19 spheroids **(Fig. 4E)**. Neither response was observed in EV controls. The reduction in IL-6 secretion following mutant p53β R273H expression was consistent with the downregulation of *ILc* by RNA-seq **(Fig. 3D)**, suggesting distinct secretory responses in WT and mutant R273H GBM cells.

Finally, induction of WT or mutant R273H p53β significantly increased *CDKN1A* (p21) and *BBC3* (*PUMA*) mRNA expression in U87-MG and SNB19 spheroids, respectively **(Fig. 4F)**, consistent with our monolayer results **(Fig. 2A,B)**.

### WT p53β reduces tumor burden and prolongs event-free survival in an orthotopic GBM xenograft model

To determine whether WT p53β suppresses GBM tumor growth *in vivo*, we intracranially injected NSG mice with U87-MG (*TP53* WT) cells expressing DOX-inducible WT p53β or an EV vector. Mice were randomized to doxycycline-supplemented (Dox) or control diets one week after injection, and tumor burden was monitored weekly by MRI **(Fig. 5A,B)**.

**Figure 5:**
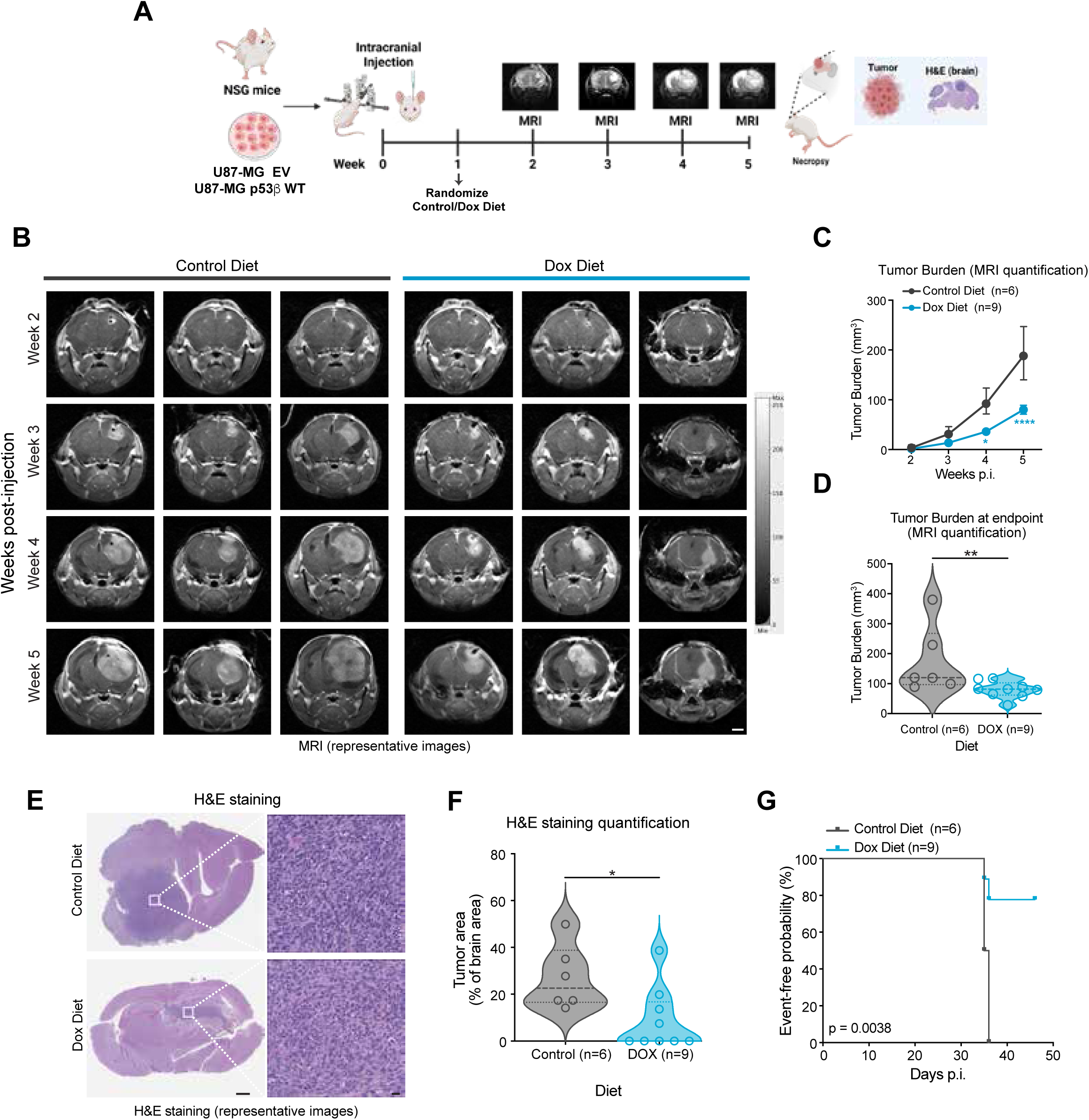
WT p53β expression suppresses tumor growth in an intracranial U87-MG GBM xenograft model. NSG mice were intracranially injected with U87-MG cells expressing doxycycline-inducible empty vector (EV) or WT p53β. Mice were randomized to a control or doxycycline-containing (DOX) diet one-week post-injection. Data shown in panels B-G correspond to mice bearing WT p53β-expressing tumors (control, n=6; DOX, n=9) unless otherwise stated. **(A)** Schematic of the experimental design and timeline. **(B)** Representative MRI images from three mice per group obtained at weeks 2 and 5 post-injection. Scale bar: 2mm. **(C)** MRI-based quantification of tumor burden (mm^3^) at the indicated times post-injection. Data are presented as mean +/- SEM. Statistical significance was assessed by two-way ANOVA followed by Sidak’s multiple comparisons test (week 2, p>0.9999; week 3, p=0.8897; week 4, p=0.0136; week 5, p<0.0001). **(D)** Quantification of tumor burden at endpoint. Mann-Whitney test, p=0.0048. **(E)** Representative hematoxylin and eosin (HCE)-stained brain sections (left) and higher-magnification images of boxed regions (right). Scale bars: 1mm (left) and 20 μm (right). **(F)** Quantification of tumor area as a percentage of total brain area in HCE-stained sections (Welch’s t-test, p=0.0284). **(G)** Kaplan-Meier curve showing exploratory event-free survival analysis, with an event defined as the first occurrence of any of the following: 10% body weight loss, tumor volume >100 mm^3^ (average at endpoint), or onset of neurological signs (hunched posture, circling, head tilt) (log-rank (Mantel-Cox) test, p=0.0038). Individual data points represent individual mice (D,F). *p < 0.05; **p<0.01; ***p<0.001; ****p<0.0001. ns, not significant.

DOX-induced WT p53β **(Supplementary Fig. 4A)** significantly reduced tumor burden compared with control-diet mice at weeks 4 and 5 post-injection (**Fig. 5C**, week 4, p=0.0136; week 5, p<0.0001), and at study endpoint (**Fig. 5D**, p=0.0048). Consistently, histological analysis revealed a significant reduction in tumor area relative to brain area (**Fig. 5E,F**, p=0.0284). WT p53β induction also reduced brain weight without significantly affecting body weight **(Supplementary Fig. 4B-E)**, and prolonged event-free survival compared with controls (**Fig. 5G**; p=0.0038). In contrast, DOX treatment of mice harboring EV-transduced U87-MG cells did not significantly affect tumor burden, histological tumor area, or event-free survival **(Supplementary Fig. 4F-O)**.

Together, these data demonstrate that WT p53β expression suppresses intracranial GBM tumor growth and prolongs event-free survival; therefore, WT p53β tumor-suppressive effects extend *in vivo*.

### Mutant p53β R273H diminishes tumor growth rate and extends event-free survival in an intracranial xenograft mouse model of glioblastoma

To determine whether mutant p53β R273H retains tumor suppressive activity *in vivo*, NSG mice were intracranially injected with U251 (*TP53* R273H) cells expressing doxycycline-inducible mutant p53β R273H or an EV control. Unlike the WT model, tumor engraftment was confirmed by MRI before mice were randomized to control (n=8) or doxycycline-supplemented (n=9) diets **(Fig. 6A,B)**. While mean pretreatment tumor volumes did not differ between groups, baseline tumor volumes varied significantly among individual animals within each group **(Supplementary Fig. 5A-C)**. Therefore, tumor growth was expressed as fold change relative to tumor volume at pretreatment per mouse.

**Figure 6:**
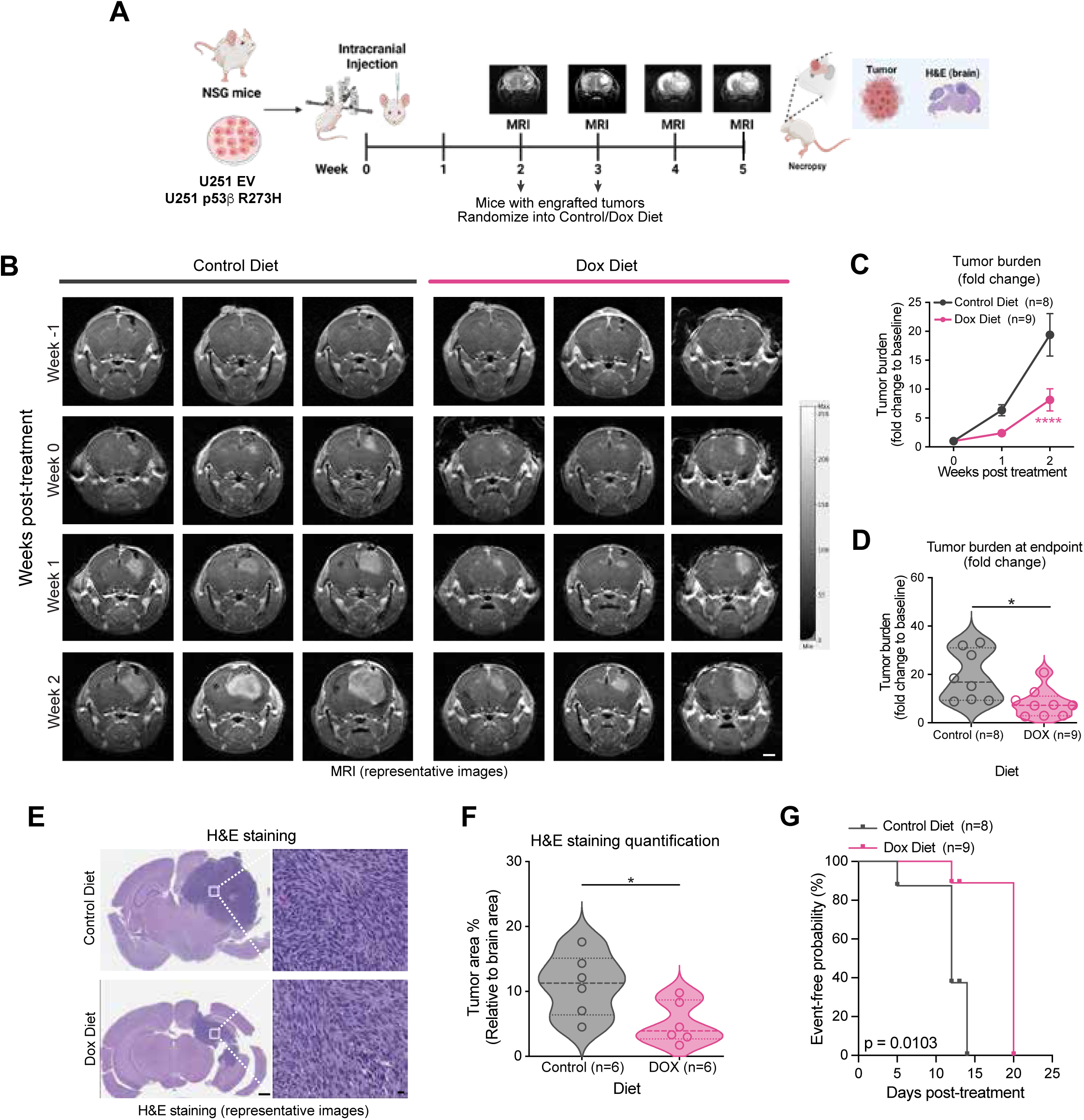
Mutant p53β R273H expression suppresses tumor growth in an intracranial U251 GBM xenograft model. NSG mice were intracranially injected with U251 cells expressing doxycycline-inducible empty vector (EV) or mutant R273H p53β. Mice were imaged by MRI starting at week 2 post-injection, and tumor-bearing mice were randomized to a control or doxycycline-containing diet (DOX) diet when a tumor was detected. Data shown in panels B-G correspond to mice bearing mutant p53β R273H-expressing tumors (control, n=8; DOX, n=9, except as indicated). Results from the EV control cohort are presented in Supplementary Figure 5. **(A)** Schematic of the experimental design and timeline. **(B)** Representative MRI images from three mice per group obtained between weeks -1 and 2 relative to treatment initiation (week 0). Scale bar = 2 mm. **(C)** MRI-based quantification of tumor burden expressed as fold change relative to each animal’s pretreatment tumor volume (baseline, week 0). Data are presented as mean +/- SEM. Statistical significance was assessed by two-way ANOVA followed by Sidak’s multiple comparisons test (Week 0, p>0.9999; Week 1, p= 0.2864; Week 2, p<0.0001). **(D)** MRI-based quantification of tumor burden (fold change relative to baseline) at endpoint (1 day before necropsy). (two-tailed Mann-Whitney test (two-tailed), p=0.0111). **(E)** Representative hematoxylin and eosin (HCE)-stained brain sections (left) and higher-magnification images of the boxed regions (right). Scale bars: 1 mm (left) and 20 µm (right). **(F)** Quantification of tumor area as a percentage of total brain area in HCE-stained sections. Only mice with a pretreatment tumor volume <3.5mm^3^ were included (control, n=6; DOX, n=6; Welch’s t-test, p=0.0436). **(G)** Kaplan-Meier curve showing exploratory event-free survival analysis, with an event defined as the first occurrence of any of the following: 10% body weight loss, tumor burden> 14-fold relative to baseline (average across groups), or onset of neurological signs (hunched posture, circling, head tilt) (log-rank (Mantel-Cox) test, p=0.0103). Individual data points represent individual mice (D,F). *p < 0.05; **p<0.01; ***p<0.001; ****p<0.0001. ns, not significant.

DOX-treatment induced mutant p53β R273H mRNA expression *in vivo* **(Supplementary Fig. 5D)** and significantly reduced tumor growth (fold change) at two weeks post-treatment (**Fig. 6C**, p<0.0001) and at endpoint (**Fig. 6D**, p=0.0111). Histological analysis restricted to animals with pretreatment tumor volumes below 3.5mm^3^ (n=6 per group) also showed significantly reduced tumor area (relative to total brain area) in mutant p53β R273H-expressing animals (**Fig. 6E, F**; p=0.0436). Although absolute and relative brain weights did not differ significantly between groups (Supplementary Fig. 5E,F), DOX-treated mice showed a significantly greater percentage body-weight change at two weeks post-treatment **(Supplementary Fig. 5G,H)**. In an exploratory event-free survival analysis, DOX-mediated induction of mutant p53β R273H was associated with a significant delay in the occurrence of study-defined disease-associated events compared with the control-diet group (**Fig. 6G**; p=0.0103).

In EV-transduced U251 tumor-bearing animals, DOX and control diets did not differ significantly in tumor growth, histological tumor area, or event-free survival **(Supplementary Fig. 5I-R)**. Together, these findings demonstrate that p53β harboring the oncogenic R273H hotspot mutation retains tumor-suppressive activity in vivo, reducing intracranial GBM growth and delaying disease-associated events.

## Discussion

Our findings demonstrate that p53β retains tumor-suppressive activity despite harboring the oncogenic *TP53* R273H hotspot mutation. Mutant p53β R273H promoted senescence, suppressed GBM proliferation, spheroid growth, invasion and intracranial tumor growth, and delayed disease-associated events in SF268, SNB19 and U251 (*TP53* R273H) GBM cells. WT p53β similarly suppressed GBM growth, consistent with its senescence-inducing activity in normal cells [12–14]. Although R273H impairs canonical p53α transcriptional activity and confers gain-of-function properties [18, 20], the same mutation remains compatible with tumor suppression in p53β, emphasizing that the functional consequences of *TP53* mutations depend, at least in part, on p53 isoform context, consistent with our previous findings [25].

Because the GBM cells used in our functional studies (U87-MG, A172, SNB19, SF268) harbor WT or R273H *TP53* gene and express endogenous p53 isoforms, including p53α, the tumor-suppressive effects of p53β overexpression should be interpreted in the context of these endogenous isoforms. Indeed, our mechanistic studies revealed functional cooperation between mutant p53β R273H and mutant p53α R273H. In *TP53*-null GBM cells, neither mutant isoform alone induced *CDKN1A* expression, whereas co-expression restored its induction. Similarly, co-expression activated the p21 promoter in a p53α-dose-dependent manner, even though neither isoform alone activated the reporter. Moreover, mutant p53β R273H co-immunoprecipitated with mutant p53α R273H in the presence of the p21 promoter-reporter construct, consistent with previous observations involving WT p53 isoforms [22]. These findings suggest that mutant p53β R273H cooperates with mutant p53α R273H to promote canonical p53 target gene expression. However, whether this association involves direct protein-protein binding or additional regulatory proteins remains unclear. Furthermore, because R273H disrupts sequence-specific DNA binding, transcriptional activation may involve alternative promoter recruitment mechanisms, altered cofactor interactions, or stabilization of promoter-associated complexes. Further studies examining endogenous promoter occupancy and associated transcriptional regulators will be needed to establish the underlying mechanism and determine its contribution to the tumor-suppressive phenotypes observed.

Transcriptomic analyses showed distinct responses to WT and mutant R273H p53β expression. WT p53β mainly induced canonical p53-associated genes and suppressed cell-cycle progression. In both mutant GBM lines, mutant p53β R273H suppressed MYC targets v1 and mTORC1 signaling, pathways that support proliferation and anabolic metabolism [26, 27]. Reduced *MYC* and *SNAI2* expression also aligned with diminished proliferation and invasion in vitro. In SF268 (L2P), HALLMARK P53 pathway enrichment and qRT-PCR-detected *CDKN1A, BAX,* and *BBC3* induction indicated partial activation of canonical p53-associated transcription. Despite substantial variation in pathway enrichment across GBM lines with the same *TP53* status, responses aligned with the anti-proliferative effects of p53β induction. This heterogeneity may reflect differences in cellular background and statistical criteria for pathway enrichment, despite accounting for doxycycline effects. Mutant R273H p53β may therefore suppress GBM growth through canonical and noncanonical transcriptional programs; the contribution of each pathway remains unclear.

Clinical association studies provide additional support for a protective role of p53β in cancers harboring mutant *TP53*. Higher p53β expression correlated with lower tumor stage and improved recurrence-free and overall survival in clear-cell renal cell carcinoma, independently of *TP53* status [28], and with reduced tumor size and improved disease-free survival, particularly in patients with mutant *TP53* [29]. In the latter study, patients with mutant *TP53* and elevated p53β expression had survival outcomes comparable to WT *TP53* patients [29]. Additional studies have implicated p53 isoform expression in cancer prognosis and treatment responses [30–34], although the direction of these associations varies by tumor type and isoform [32–34]. Importantly, these correlative studies did not establish causality or mutant p53β function; our findings provide functional evidence of its tumor-suppressive activity in GBM.

Beyond proliferation, p53β expression also suppressed spheroid growth and invasion. Reduced N-cadherin expression, EMT-associated transcriptional changes, and *SNAI2* downregulation suggest that p53β modulates invasion-associated pathways. WT p53β increased IL-6 secretion in U87-MG spheroids, whereas mutant R273H p53β decreased IL-6 secretion in SNB19 spheroids, consistent with IL6 downregulation by RNA-seq. In this context, senescence-associated inflammatory signaling can suppress tumor growth, or, through sustained SASP signaling, promote tumor progression and therapy resistance [35, 36]. However, IL-6 alone does not define the SASP, and differences between cell lines preclude attributing these responses exclusively to *TP53* status. The consequences for the tumor microenvironment and recurrence warrant further investigation.

Our findings also identify SRSF3-mediated regulation of p53β expression as a potential therapeutic target in GBM. We previously demonstrated that SRSF3 negatively regulates p53β alternative splicing and that its inhibition promotes cellular senescence in normal human cells [12, 13, 15]. Here, SRSF3 depletion increased p53β expression and induced senescence- and apoptosis-associated responses in GBM cells harboring either WT or R273H mutant *TP53*. The absence of senescence and p53 target induction in *TP53-*null cells supports the dependence of these responses on *TP53* expression. However, because SRSF3 regulates numerous transcripts beyond *TP53,* the specific contribution of p53β induction to the effects of SRSF3 depletion remains to be determined.

The therapeutic potential of p53β regulation is further supported by studies showing that pharmacological upregulation of p53β and p53γ through inhibition of nonsense-mediated mRNA decay can activate p53-associated pathways and suppress tumor growth [37]. Nevertheless, systemic SRSF3 inhibition may affect normal tissues, including astrocytes, in which SRSF3 depletion also induces senescence [14]. Tumor-selective approaches, including targeted delivery of SRSF3 siRNA [38] or more selective modulation of p53β alternative splicing, warrant further investigation. Several limitations of this study should be considered. Our functional studies relied primarily on inducible p53β expression, and whether restoring endogenous p53β to physiological levels produces comparable effects remains to be determined. In addition, although full-length p53β was the predominant isoform detected, the contribution of other isoforms detected at lower levels, potentially corresponding to Δ40p53β, Δ133p53β and Δ160p53β, to the observed tumor-suppressive phenotypes cannot be excluded. Furthermore, WT and mutant R273H p53β were examined in different cellular backgrounds, which may have influenced the observed responses. Our intracranial studies used GBM xenografts in immunodeficient NSG mice, limiting assessment of the effects of p53β expression on antitumor immunity and the tumor microenvironment. Finally, our findings are restricted to the R273H hotspot mutation, and whether other oncogenic *TP53* mutations retain tumor-suppressive activity in the p53β isoform remains to be established [39].

In summary, mutant p53β R273H isoform retains tumor-suppressive activity in GBM cells harboring *TP53* R273H and cooperates with p53α R273H to partially restore canonical p53-associated transcription and supress growth- and invasion-promoting pathways. These findings emphasize the importance of isoform context in the functional consequences of *TP53* mutations and support further investigation of p53β restoration as a therapeutic strategy in GBM.

## Methods

### TCGA analysis

*SRSF3* mRNA expression, *TP53* mutational status, and clinical data were obtained from TCGA via cBioPortal, the TCGAbiolinks, and the Genomics Data Commons (516 lower-grade glioma, 154 GBM, and five normal brain samples before reclassification). Gliomas were reclassified by *IDH* status using 2021 WHO criteria; SRSF3 expression was compared across tumor classifications and *TP53* status.

### Cell lines and cell culture

Human GBM cell lines are listed in Supplementary Table S1. Certificates of analysis were available from their sources, and cells underwent MTBM testing, including Mycoplasma testing, upon receipt. Cells were cultured in our laboratory for fewer than 20 consecutive passages and were maintained in DMEM supplemented with 10% FBS, 1% penicillin/streptomycin solution, and 1% L-glutamine.

### siRNA transfection

SRSF3 was depleted using specific siRNAs (15nM) using Lipofectamine RNAiMAX (ThermoFisher) per the manufacturer’s instructions.

### Doxycycline-inducible p53β expression vectors

pLV lentiviral constructs and viral particles are listed in Supplementary Table S1. Cells were transduced with lentiviral Tet3G (MOI 3), selected with puromycin (1 μg/ml), and subsequently transduced with empty vector (EV), Flag-tagged WT, or mutant R273H p53β (MOI 1). Cells were selected with blasticidin (1 μg/ml) and treated with doxycycline (DOX, 0.1 μg/ml) or vehicle (DMSO).

### Senescence-associated β-galactosidase staining

SA-β-gal staining was performed with the Senescence Associated (SA)-β-Galactosidase Staining Kit (Cell Signaling Technology, Danvers, MA, USA).

### Cell confluence imaging

Cell proliferation was monitored in 96-well plates (5×10^2^ cells/well) by Incucyte-S3® (10x magnification) every 6h following DOX (0.1-1μg/ml) or DMSO treatment.

### Immunofluorescence

Immunofluorescence staining for p21, Ki67, and cleaved caspase-3 was performed as previously described [40]. At least 3 fields of view and a total of 50-100 cells were quantified per slide. Antibodies used are listed in Supplementary Table S1.

### Western Blot

Western blot was performed as described previously [25]. Antibodies used are listed in Supplementary Table S1. Full-size western blot images are in Supplementary Fig. 6.

### Quantitative Real-Time Polymerase Chain Reaction (qRT-PCR)

qRT-PCR was performed as previously described [41]. Expression was normalized to GAPDH using the ΔΔCt method. Commercial kits, qPCR assays, and sequences are listed in Supplementary Table S1.

### Transcriptomic Analysis

U87-MG, A172, SF268 and SNB19 cells stably expressing doxycycline-inducible (DOX, Tet-On) the corresponding WT or mutant R273H p53β, or an empty control, were harvested 48 hours after treatment with DOX or a DMSO control (three biological replicates per condition). RNA samples (RIN > 8) underwent stranded mRNA library preparation and paired-end 150-bp sequencing on Illumina NovaSeq (Genewiz/Azenta). Reads were trimmed using Trimmomatic v.0.36 and aligned to the human GRCh38 reference genome using the STAR aligner v.2.5.2b.

Differential gene expression and pathway analyses used NIH Integrated Data Analysis Platform (NIDAP 1.0) as previously described [25]. DOX versus DMSO contrasts were analyzed separately per cell line and vector (DEGs: |log_2_FC| > 1, adjusted P<0.05). *TP53* was excluded because reads from the induced p53β transgene could not be reliably distinguished from endogenous *TP53*. GSEA was performed per cell line using t-statistic-ranked genes and Hallmark, KEGG, Reactome, and GO gene sets (Benjamini–Hochberg adjusted *P* < 0.05). Pathways significantly enriched in matched EV DOX vs DMSO controls were excluded from the corresponding p53β result. L2P used the Hallmark gene sets. GEO accession number GSE355830.

### Dual luciferase assay

U87-MG and U251 cells expressing inducible WT or mutant R273H p53β (or EV) were seeded, treated overnight with DMSO or doxycycline, and co-trasfected with Renilla, p21 promoter-firefly luciferase, and p53α (0, 1, 3.3, and 10 ng) [22]. Luciferase activity was measured with a Dual-Luciferase Reporter Assay Kit (Promega) following the manufacturer’s instructions, and firefly activity was normalized to Renilla.

### Co-immunoprecipitation

Doxycycline-inducible Flag-p53β (WT or R273H) cells were transfected with pcDNA3 control or WT/R273H p53α (2 μg), with or without p21-luc (2 μg). Protein lysates were immunoprecipitated using anti-Flag M2 affinity gel according to the manufacturer’s instructions and analyzed by immunoblotting. Reagent details are provided in Supplementary Table S1.

### 3D spheroid growth and invasion assays

GBM spheroids were established in Tumorsphere MediumX and treated after 48h incubation with DOX (0.1 μg/ml) or DMSO. Spheroid growth and invasion experiments were performed and analyzed as previously described[25].

### IL-6 ELISA

Spheroid supernatants were harvested at day 7 post-incubation. IL-6 secretion was measured using the Human IL-6 ELISA kit (Sigma-Aldrich) following the manufacturer’s instructions.

### Orthotopic GBM xenograft mouse model

#### General procedures

Animal procedures were approved by the NCI Frederick Animal Care and Use Committee (ASP #22-334) and followed the ACUC (https://ncifrederick.cancer.gov/Lasp/Acuc/) and ARRIVE (https://arriveguidelines.org/) guidelines. Female NSG mice (6-8 weeks old) received stereotaxic intracranial injections of 2×10^5^ cells in 4 μl PBS (3 mm posterior, 2 mm right of bregma, at a depth of 2 mm below the dura) [42]. Body weight and neurological signs were monitored at least twice weekly. Humane endpoints included 20% body-weight loss, tumor volume >200 mm³, or neurological signs. Allocation, exclusions, and group sizes are detailed in Supplementary Table S2.

#### U87-MG mouse model

Surviving mice were randomized to control or DOX diets one week after intracranial injection. Tumor burden was monitored weekly by MRI from week 2 post-injection (Fig. 5A).

#### U251 mouse model

U251 was selected for its established tumorigenicity and GBM histopathology [43]. Mice with MRI-confirmed engraftment at 2-3 weeks post-injection were randomized to control or DOX diets one day after imaging. Tumor burden was monitored weekly and expressed as fold change relative to pretreatment MRI volume (Fig. 6A).

#### Disease-associated event-free survival analysis

Because studies ended after scheduled MRI assessments, we defined an event as the first occurrence of neurological signs (hunched posture, circling, or head tilt), ≥10% body-weight loss, or an MRI-based tumor burden threshold of >100 mm³ (U87-MG) or >14-fold increase in pretreatment volume (U251). Both tumor thresholds were selected retrospectively from the observed endpoint distributions and correspond to the mean across groups per animal study. Event-free probability was estimated using Kaplan–Meier analysis and compared by log-rank test.

### Magnetic resonance imaging (MRI)

MRI was performed using a 3.0 T Philips Ingenia Elition X scanner with T2-weighted and pre/post-contrast T1-weighted sequences (gadopiclenol, 0.05 mmol/kg). During anesthesia with 1.5-2% isoflurane, respiration and body temperature (maintained at 36–37 °C) were continuously monitored. Tumor volume was quantified from post-contrast T1-weighted images using ITK-SNAP software.

### Histological analysis

Brains were fixed in formalin, paraffin-embedded, and sectioned sagittally at 5 µm for hematoxylin and eosin (HCE) staining. Tumor burden was quantified and expressed as a percentage of total brain area, and slides were assessed by a blinded, board-certified veterinary pathologist. Because baseline histology was unavailable, U251 analysis was restricted to mice with pretreatment MRI tumor burden <3.5 mm^3^ (mean baseline tumor burden across groups).

### Statistical analyses

At least three independent experiments were performed for in vitro assays. Statistical tests are specified in the figure legends. *P* <0.05 was considered significant.

## Ethics statement and consent to participate

Publicly available TCGA data required no additional human-subject approval.

## Acknowledgements

We thank Dr. Orieta Celiku and Dr. Mark Gilbert (Neuro Oncology Branch, NCI/NIH) for assistance with TCGA data analysis; Jessica Ebersole and the Animal Research Technical Support (Frederick National Laboratory of Cancer Research, NIH) for animal care and support; Dr. Claudia Mani and the Small Animal Research Program (SAIP/NCI/NIH) team for MRI support; Dr. Elijah Edmonson and the Molecular Histopathology Laboratory (MHL/NCI/NIH) for necropsy and histological analysis; Dr. Luowei Li for Incucyte support. We also thank the CCBR team (NCI/NIH) for their assistance with RNA-sequencing data analysis performed in the NIH Integrated Data Analysis Platform (NIDAP). Venn diagrams were generated using https://www.interactivenn.net/. Figures 1A, 5A, and 6A were created using BioRender.com.

This research was supported by the Intramural Research Program of the National Institutes of Health (NIH) (ZIA BC 011496). Contributions from NIH-affiliated authors were made in the course of their official duties as federal employees and are considered works of the United States Government in accordance with agency policy requirements. The findings and conclusions presented herein are solely those of the authors and do not necessarily represent the official views of the NIH or the U.S. Department of Health and Human Services.

## Authors Contributions

NVM conceived and designed the study, developed animal study protocols and ensured compliance with ACUC requirements and ARRIVE guidelines, performed in vitro experiments, analyzed and interpreted data, and was responsible for manuscript preparation and submission. JAB performed the SRSF3-related in vitro experiments and analyzed and interpreted the corresponding data (Supplementary Fig. 1), with technical support from KU and AT. CZ assisted with MRI-based tumor quantification. SN and SJ provided experimental expertise and technical support for Western blotting, while SJ also assisted with dual-luciferase reporter assays, co-immunoprecipitation, and spheroid assays. CCH and IH provided scientific input on study design, data interpretation, and manuscript content. All authors reviewed and approved the final manuscript.

## References

1 Ostrom QT, Gittleman H, Truitt G, Boscia A, Kruchko C, Barnholtz-Sloan JS. CBTRUS Statistical Report: Primary Brain and Other Central Nervous System Tumors Diagnosed in the United States in 2011-2015. Neuro Oncol 2018; 20: iv1–iv86.

2 Brown NF, Ottaviani D, Tazare J, Gregson J, Kitchen N, Brandner S et al. Survival Outcomes and Prognostic Factors in Glioblastoma. Cancers (Basel) 2022; 14.

3 Mendiratta G, Ke E, Aziz M, Liarakos D, Tong M, Stites EC. Cancer gene mutation frequencies for the U.S. population. Nat Commun 2021; 12: 5961.

4 Zhang Y, Dube C, Gibert M, Jr., Cruickshanks N, Wang B, Coughlan M et al. The p53 Pathway in Glioblastoma. Cancers (Basel) 2018; 10.

5 Xiong Y, Zhang Y, Xiong S, Williams-Villalobo AE. A Glance of p53 Functions in Brain Development, Neural Stem Cells, and Brain Cancer. Biology (Basel) 2020; 9.

6 Manfredi JJ. Inactivation of Wild-Type p53 by Asparagine Endopeptidase in Glioblastoma: An Opportunity to Target the “Undruggable”. J Natl Cancer Inst 2020; 112: 327–329.

7 Lin Y, Liao K, Miao Y, Qian Z, Fang Z, Yang X et al. Role of Asparagine Endopeptidase in Mediating Wild-Type p53 Inactivation of Glioblastoma. J Natl Cancer Inst 2020; 112: 343–355.

8 Kraus A, Gross MW, Knuechel R, Munkel K, Neff F, Schlegel J. Aberrant p21 regulation in radioresistant primary glioblastoma multiforme cells bearing wild-type p53. J Neurosurg 2000; 93: 863–872.

9 Beck J, Turnquist C, Horikawa I, Harris C. Targeting cellular senescence in cancer and aging: roles of p53 and its isoforms. Carcinogenesis 2020; 41: 1017–1029.

10 Bourdon JC, Fernandes K, Murray-Zmijewski F, Liu G, Diot A, Xirodimas DP et al. p53 isoforms can regulate p53 transcriptional activity. Genes Dev 2005; 19: 2122–2137.

11 Joruiz SM, Bourdon JC. p53 Isoforms: Key Regulators of the Cell Fate Decision. Cold Spring Harb Perspect Med 2016; 6.

12 Fujita K, Mondal AM, Horikawa I, Nguyen GH, Kumamoto K, Sohn JJ et al. p53 isoforms Delta133p53 and p53beta are endogenous regulators of replicative cellular senescence. Nat Cell Biol 2009; 11: 1135–1142.

13 Mondal AM, Horikawa I, Pine SR, Fujita K, Morgan KM, Vera E et al. p53 isoforms regulate aging- and tumor-associated replicative senescence in T lymphocytes. J Clin Invest 2013; 123: 5247–5257.

14 Turnquist C, Horikawa I, Foran E, Major EO, Vojtesek B, Lane DP et al. p53 isoforms regulate astrocyte-mediated neuroprotection and neurodegeneration. Cell Death Differ 2016; 23: 1515–1528.

15 Tang Y, Horikawa I, Ajiro M, Robles AI, Fujita K, Mondal AM et al. Downregulation of splicing factor SRSF3 induces p53beta, an alternatively spliced isoform of p53 that promotes cellular senescence. Oncogene 2013; 32: 2792–2798.

16 Song X, Wan X, Huang T, Zeng C, Sastry N, Wu B et al. SRSF3-Regulated RNA Alternative Splicing Promotes Glioblastoma Tumorigenicity by Affecting Multiple Cellular Processes. Cancer Res 2019; 79: 5288–5301.

17 Fuentes-Fayos AC, Vazquez-Borrego MC, Jimenez-Vacas JM, Bejarano L, Pedraza-Arevalo S, F LL et al. Splicing machinery dysregulation drives glioblastoma development/aggressiveness: oncogenic role of SRSF3. Brain 2020; 143: 3273–3293.

18 Cho Y, Gorina S, Jeffrey PD, Pavletich NP. Crystal structure of a p53 tumor suppressor-DNA complex: understanding tumorigenic mutations. Science 1994; 265: 346–355.

19 Dittmer D, Pati S, Zambetti G, Chu S, Teresky AK, Moore M et al. Gain of function mutations in p53. Nat Genet 1993; 4: 42–46.

20 Alvarado-Ortiz E, de la Cruz-Lopez KG, Becerril-Rico J, Sarabia-Sanchez MA, Ortiz-Sanchez E, Garcia-Carranca A. Mutant p53 Gain-of-Function: Role in Cancer Development, Progression, and Therapeutic Approaches. Front Cell Dev Biol 2020; 8: 607670.

21 Louis DN, Perry A, Wesseling P, Brat DJ, Cree IA, Figarella-Branger D et al. The 2021 WHO Classification of Tumors of the Central Nervous System: a summary. Neuro Oncol 2021; 23: 1231–1251.

22 Marcel V, Fernandes K, Terrier O, Lane DP, Bourdon JC. Modulation of p53beta and p53gamma expression by regulating the alternative splicing of *TP53* gene modifies cellular response. Cell Death Differ 2014; 21: 1377–1387.

23 Jiang X, Tan J, Li J, Kivimae S, Yang X, Zhuang L et al. DACT3 is an epigenetic regulator of Wnt/beta-catenin signaling in colorectal cancer and is a therapeutic target of histone modifications. Cancer Cell 2008; 13: 529–541.

24 Zhang N, Wei P, Gong A, Chiu WT, Lee HT, Colman H et al. FoxM1 promotes beta-catenin nuclear localization and controls Wnt target-gene expression and glioma tumorigenesis. Cancer Cell 2011; 20: 427–442.

25 Joruiz SM, Von Muhlinen N, Horikawa I, Gilbert MR, Harris CC. Distinct functions of wild-type and R273H mutant Delta133p53alpha differentially regulate glioblastoma aggressiveness and therapy-induced senescence. Cell Death Dis 2024; 15: 454.

26 Masui K, Tanaka K, Akhavan D, Babic I, Gini B, Matsutani T et al. mTOR complex 2 controls glycolytic metabolism in glioblastoma through FoxO acetylation and upregulation of c-Myc. Cell Metab 2013; 18: 726–739.

27 Carpenter AB, Sacknovitz A, Hanft S, Gandhi CD, Jhanwar-Uniyal M. Complex role of mTOR signaling pathway in glioblastoma and its stem cells. Adv Biol Regul 2026; 100: 101143.

28 Zhang H, Zhao Y, Sun P, Zhao M, Su Z, Jin X et al. p53beta: a new prognostic marker for patients with clear-cell renal cell carcinoma from 5.3 years of median follow-up. Carcinogenesis 2018; 39: 368–374.

29 Avery-Kiejda KA, Morten B, Wong-Brown MW, Mathe A, Scott RJ. The relative mRNA expression of p53 isoforms in breast cancer is associated with clinical features and outcome. Carcinogenesis 2014; 35: 586–596.

30 Anensen N, Oyan AM, Bourdon JC, Kalland KH, Bruserud O, Gjertsen BT. A distinct p53 protein isoform signature reflects the onset of induction chemotherapy for acute myeloid leukemia. Clin Cancer Res 2006; 12: 3985–3992.

31 Chen J, Zhang D, Qin X, Owzar K, McCann JJ, Kastan MB. DNA-Damage-Induced Alternative Splicing of p53. Cancers (Basel) 2021; 13.

32 Hofstetter G, Berger A, Fiegl H, Slade N, Zoric A, Holzer B et al. Alternative splicing of p53 and p73: the novel p53 splice variant p53delta is an independent prognostic marker in ovarian cancer. Oncogene 2010; 29: 1997–2004.

33 Groen K, Steffens Reinhardt L, Bourdon JC, Avery-Kiejda KA. It is not all about the alpha: elevated expression of p53beta variants is associated with lower probability of survival in a retrospective melanoma cohort. Cancer Cell Int 2023; 23: 228.

34 Rojas EA, Corchete LA, De Ramon C, Krzeminski P, Quwaider D, Garcia-Sanz R et al. Expression of p53 protein isoforms predicts survival in patients with multiple myeloma. Am J Hematol 2022; 97: 700–710.

35 Aasland D, Gotzinger L, Hauck L, Berte N, Meyer J, Effenberger M et al. Temozolomide Induces Senescence and Repression of DNA Repair Pathways in Glioblastoma Cells via Activation of ATR-CHK1, p21, and NF-kappaB. Cancer Res 2019; 79: 99–113.

36 Demaria M, O’Leary MN, Chang J, Shao L, Liu S, Alimirah F et al. Cellular Senescence Promotes Adverse Effects of Chemotherapy and Cancer Relapse. Cancer Discov 2017; 7: 165–176.

37 Gudikote JP, Cascone T, Poteete A, Sitthideatphaiboon P, Wu Q, Morikawa N et al. Inhibition of nonsense-mediated decay rescues p53beta/gamma isoform expression and activates the p53 pathway in MDM2-overexpressing and select p53-mutant cancers. J Biol Chem 2021; 297: 101163.

38 Nakamichi S, von Muhlinen N, Yamada L, Melamed JR, Papp TE, Parhiz H et al. SRSF3 knockdown-induced cellular senescence as a possible therapeutic strategy for non-small cell lung cancer. Carcinogenesis 2025; 46.

39 Oren M, Rotter V. Mutant p53 gain-of-function in cancer. Cold Spring Harb Perspect Biol 2010; 2: a001107.

40 Turnquist C, Beck JA, Horikawa I, Obiorah IE, Von Muhlinen N, Vojtesek B et al. Radiation-induced astrocyte senescence is rescued by Delta133p53. Neuro Oncol 2019; 21: 474–485.

41 Ungerleider K, Beck JA, Lissa D, Joruiz S, Horikawa I, Harris CC. Delta133p53alpha Protects Human Astrocytes from Amyloid-beta Induced Senescence and Neurotoxicity. Neuroscience 2022; 498: 190–202.

42 El Meskini R, Atkinson D, Weaver Ohler Z. Translational Orthotopic Models of Glioblastoma Multiforme. J Vis Exp 2023.

43 Radaelli E, Ceruti R, Patton V, Russo M, Degrassi A, Croci V et al. Immunohistopathological and neuroimaging characterization of murine orthotopic xenograft models of glioblastoma multiforme recapitulating the most salient features of human disease. Histol Histopathol 2009; 24: 879–891.

