## Supplemental Materials for "p53β, an alternatively spliced isoform of p53, induces tumor-suppressive cellular senescence and suppresses tumor growth in wild-type and mutant R273H *TP53* glioblastoma"

Supplementary Figure 1

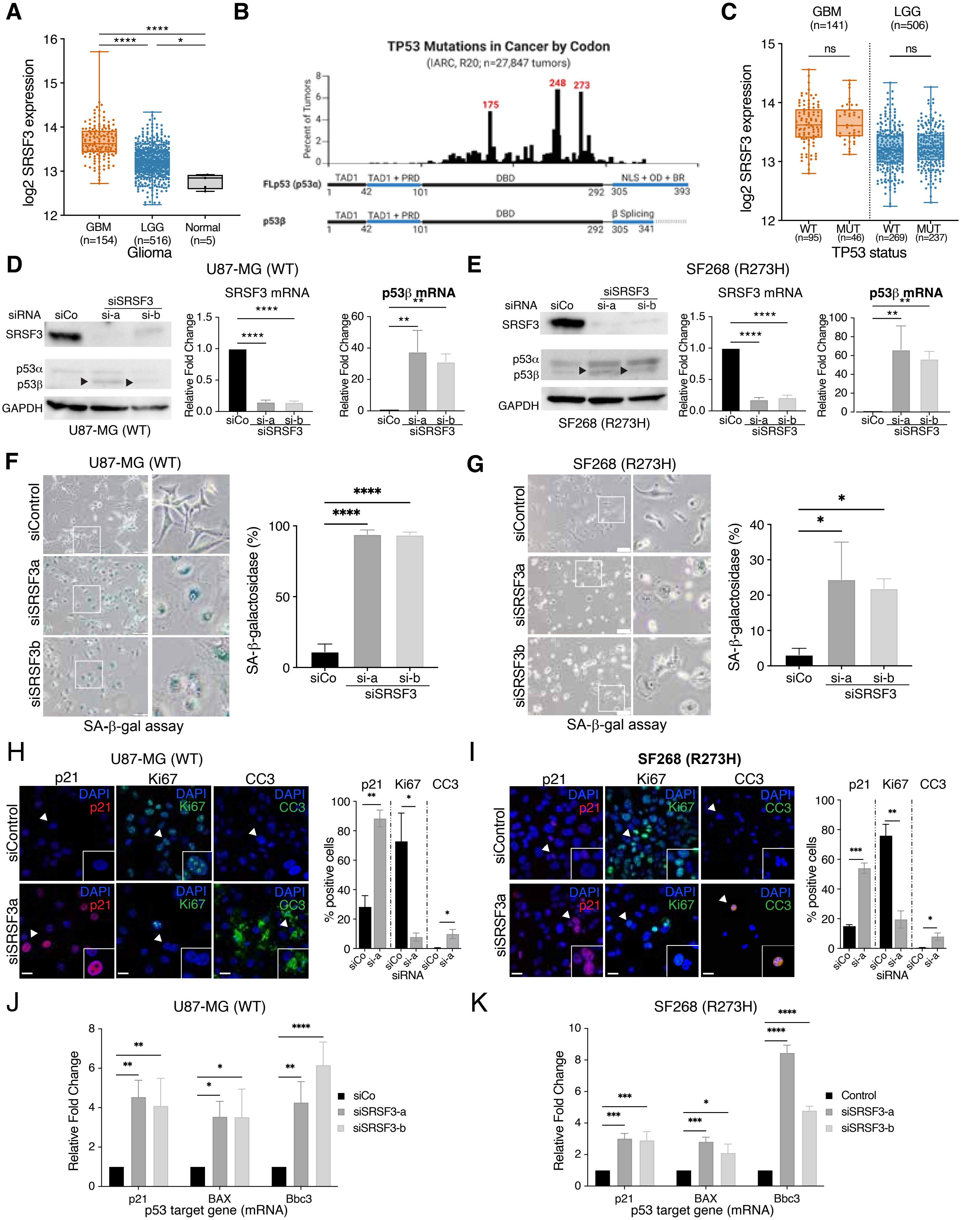

### Supplementary Figure 1

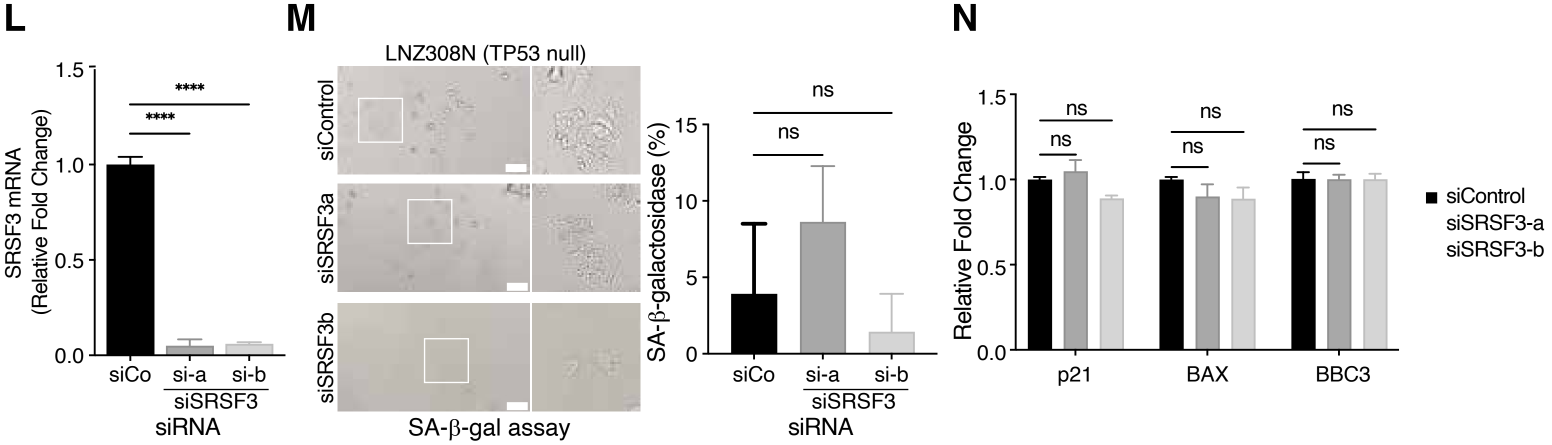

__Supplementary Figure 1__: **SRSF3 depletion restores p53β expression and induces senescence and apoptosis in GBM cells harboring WT or mutant R273H *TP53*.** **(A)** $Log_2$-transformed *SRSF3* expression in glioblastoma (GBM, N=154), low-grade glioma (LGG, N=516), and normal brain tissue (normal, N=5) samples from TCGA. **(B)** Distribution of *TP53* mutations by codon in the IARC *TP53* database (release 2, n=27,847 tumors), alongside schematic representations of the domain structure of full-length p53α and p53β. The major mutation hotspots at codons 175, 248, and 273 lie within the DNA-binding domain (DBD) shared by both isoforms. **(C)** $Log_2$-transformed *SRSF3* mRNA expression in GBM (n=141) and LGG (n=506) stratified by *TP53* mutation status. Samples lacking *TP53* mutational data were not included. **(D-K)** U87-MG (*TP53* WT) and SF268 (*TP53* R273H) GBM cells were transfected with control siRNA (siCo) or the indicated SRSF3-targeting siRNAs (siSRSF3a, siSRSF3b) and harvested four days after transfection. **(D,E)** Left: Western blot analysis of SRSF3, p53α and p53β isoforms, and GAPDH as a loading control. Arrowheads indicate p53β. Cropped blots are shown; uncropped blots are provided in Supplementary Figure 6. Right: qRT-PCR analysis of relative *SRSF3* and *p53β* mRNA expression in **(D)** U87-MG and **(E)** SF268 GBM cells transfected with the indicated siRNAs. **(F,G)** Representative images and quantification of senescence-associated β-galactosidase (SA-β-gal) activity following SRSF3 knockdown in **(F)** U87-MG and **(G)** SF268. Scale bar: 50 µm. **(H,I)** Representative immunofluorescence images and quantification of the percentage of cells positive for p21 (red), Ki67 (green), and cleaved caspase 3 (CC3; green) in **(H)** U87-MG and **(I)** SF268 GBM cells transfected with the indicated siRNAs. Nuclei were counterstained with DAPI (blue). Scale bar: 20 µm. **(J,K)** Relative mRNA expression of the p53 target genes *CDKN1A* (p21), *BAX*, and *BBC3* (*PUMA)* following SRSF3 knockdown in **(J)** U87-MG and **(K)** SF268 GBM cells. **(L-N)** *TP53*-null LNZ308N GBM cells transfected with two independent siRNAs targeting SRSF3 or a control (siCo). **(L)** qRT-PCR analysis of relative *SRSF3* mRNA expression. **(M)** Representative images and quantification of SA-β-gal activity. Scale bar: 20 µm. **(N)** Relative mRNA expression of the indicated p53 target genes. All mRNA expression measurements were normalized to *GAPDH* and expressed relative to siCo. Quantitative data from in vitro assays are presented as mean ± SD from three independent experiments (n=3). Statistical significance was assessed using Kruskal-Wallis test followed by Dunn's multiple comparisons test **(A, C)**, ordinary one-way ANOVA followed by Tukey's multiple comparisons test **(D-G, L, M)**, unpaired t-tests **(H, I)**, or two-way ANOVA followed by Tukey's multiple comparisons test **(J, K, N)**. *p < 0.05; **p<0.01; ***p<0.001; ****p<0.0001. ns, not significant.

#### Supplementary Figure 2

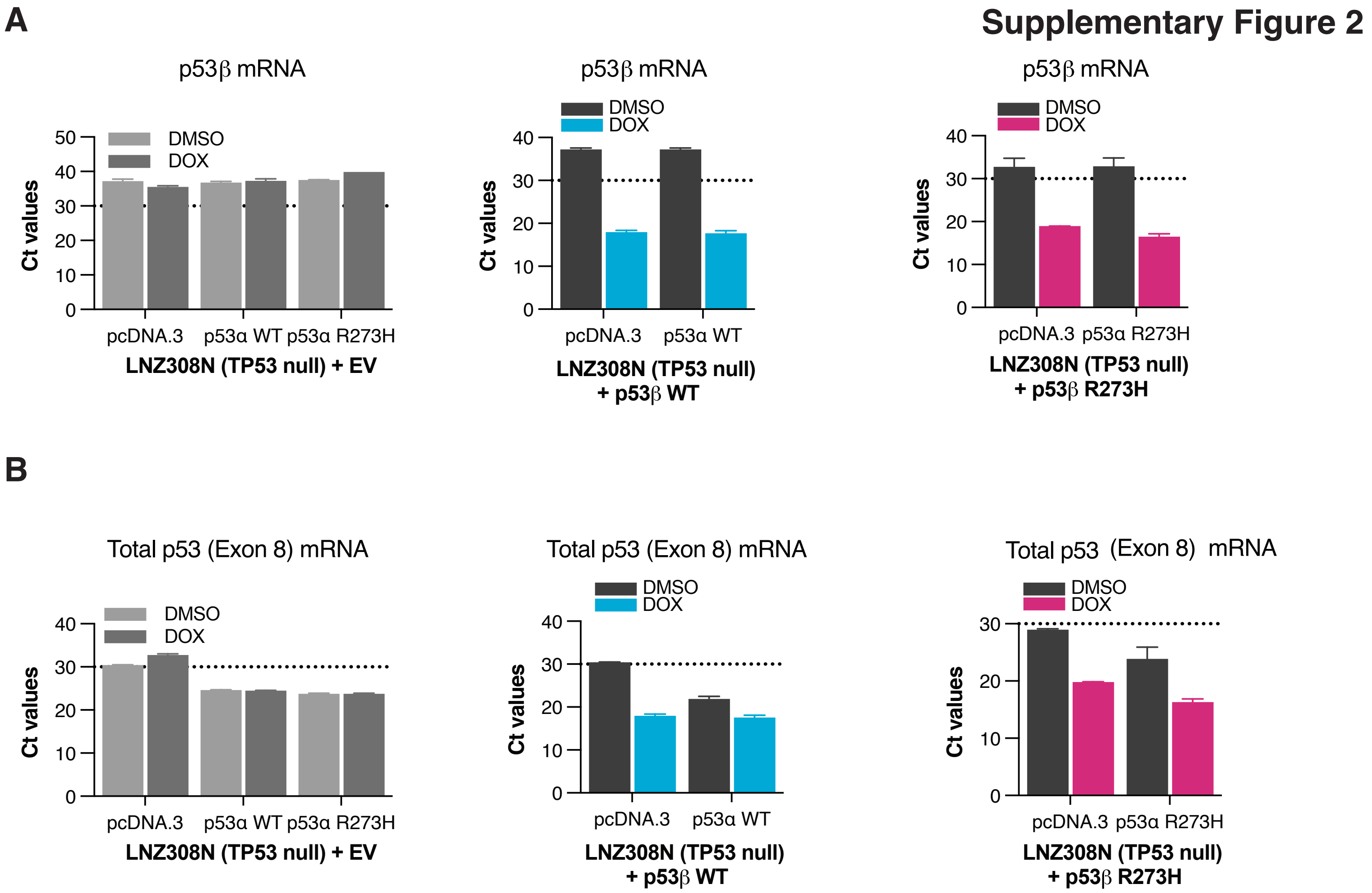

**Supplementary Figure 2: Confirmation of p53α and p53β transgene expression in *TP53*-null LNZ308N GBM cells.** Cells stably expressing the indicated doxycycline-inducible vectors (EV, WT or mutant R273H p53β) were transiently transfected with WT or mutant R273H p53α, or pcDNA.3 as a negative control, and treated with DMSO or DOX (0.1 μg/ml) for 96 h. **(A,B)** qRT-PCR analysis showing cycle threshold (Ct) values of **(A)** p53β or **(B)** Total p53 (Exon 8) mRNA. Dotted lines indicate a Ct value of 30, above which mRNA expression was considered below the reliable quantification range for these assays. Raw Ct values are presented to confirm transgene expression without calculating relative mRNA expression or performing statistical comparisons. Data are presented as mean ± SD from three independent experiments (n=3).

### Supplementary Figure 3

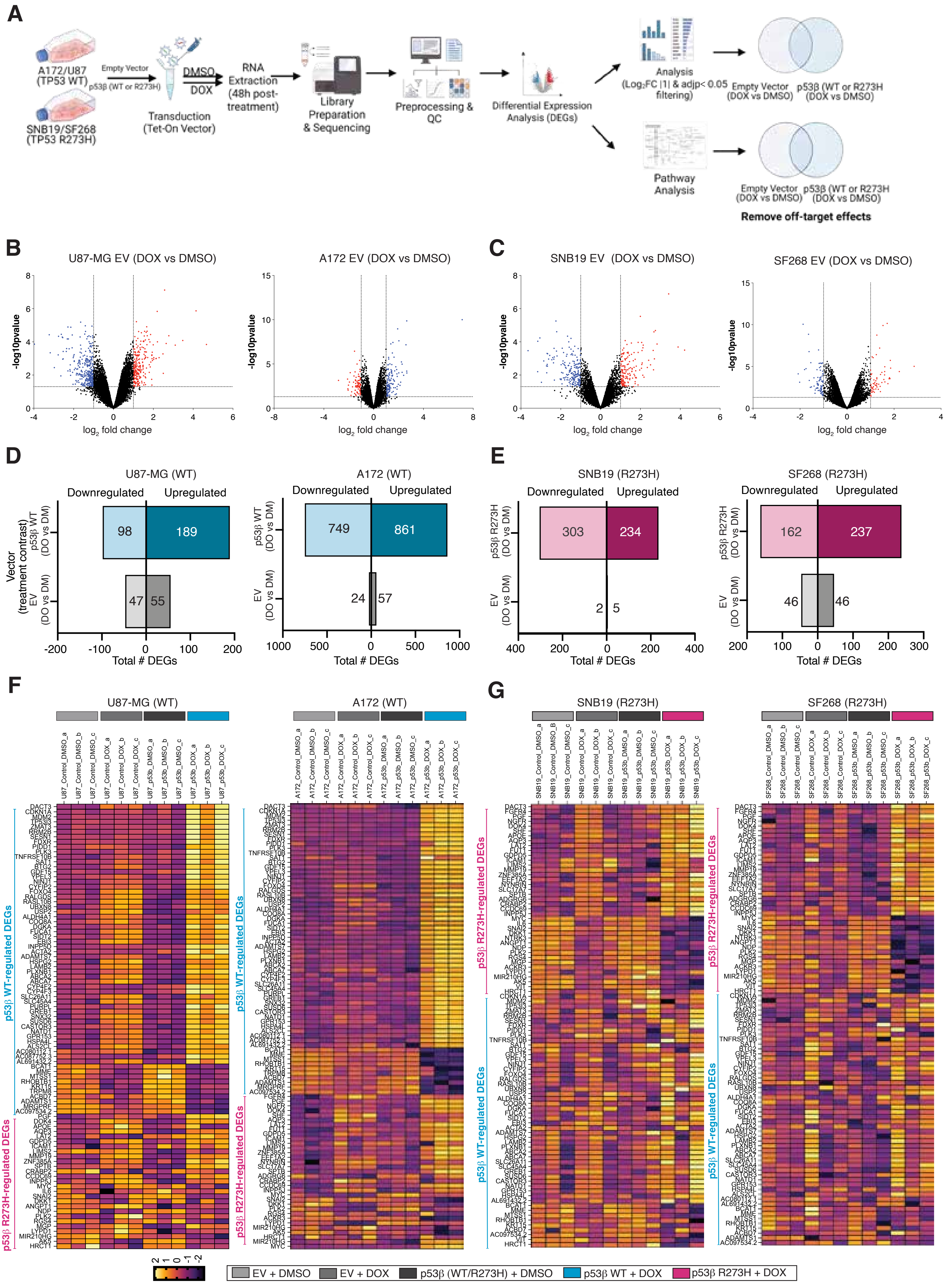

### Supplementary Figure 3

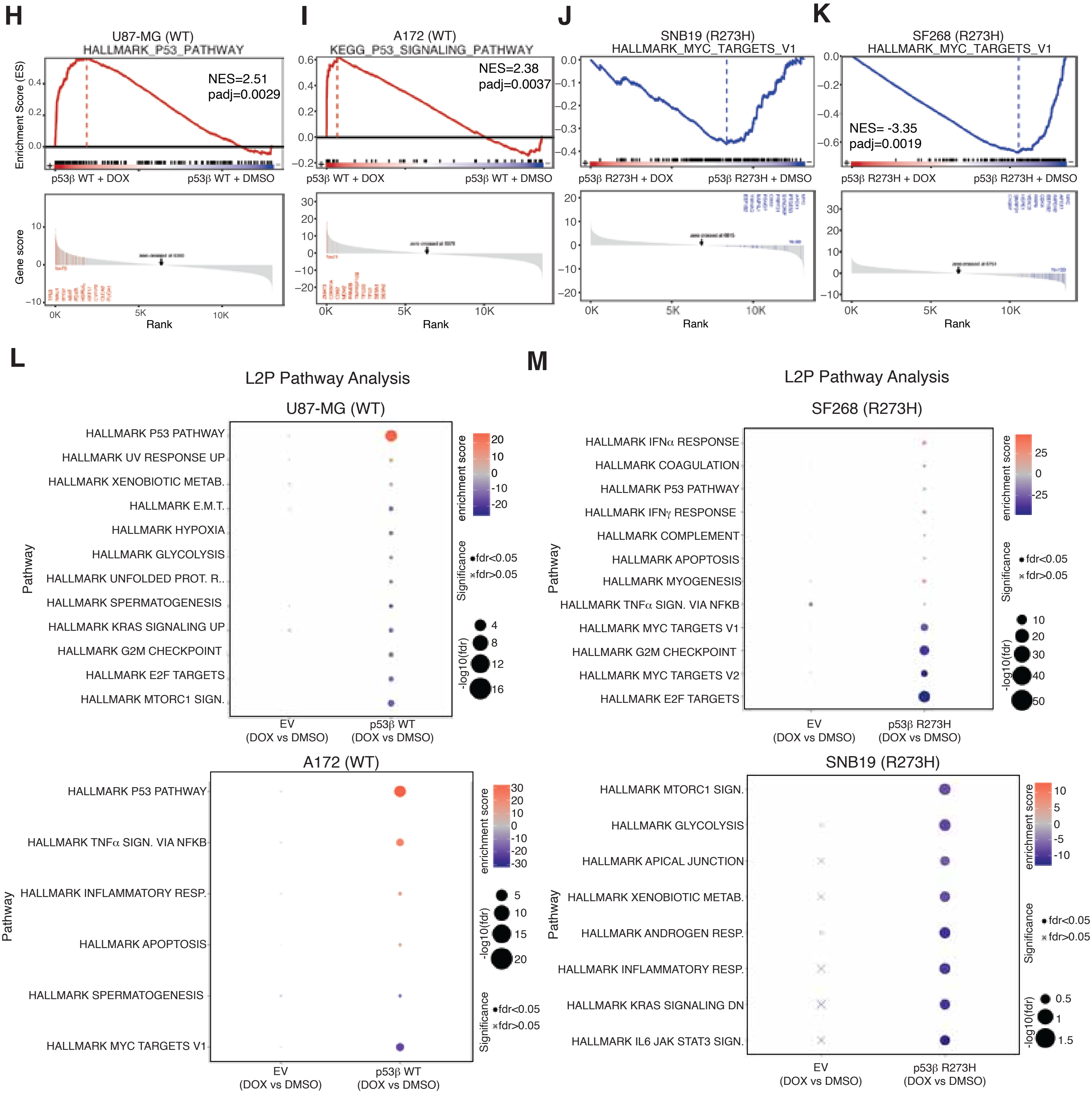

__Supplementary Figure 3:__ **RNA-seq analysis of gene expression and signaling pathways regulated by WT and mutant R273H p53β in GBM cells. (A)** Schematic of RNA-seq workflow, including doxycycline (DOX) induction, RNA extraction, sequencing, differential expression analysis, and pathway analysis. DEGs and enriched gene sets identified in the corresponding empty vector (EV) DOX vs DMSO comparisons were excluded from p53β-specific analyses to account for doxycycline-associated effects. **(B,C)** Volcano plots showing $log_2$ fold change versus -$log_{10}$ (p value) in the indicated WT **(B)** or mutant R273H **(C)** *TP53* GBM cell lines expressing doxycycline-inducible EV (DOX vs DMSO). Red and blue dots indicate significantly upregulated and downregulated genes, respectively. **(D,E)** Total number of upregulated and downregulated differentially expressed genes (DEGs) identified in the corresponding p53β and EV DOX vs DMSO contrasts in **(D)** WT or **(E)** mutant R273H GBM cell lines. DEGs were defined by $|log_2fc|>1$ and adjusted pval <0.05. **(F,G)** Heatmaps of normalized gene expression across three biological replicates per condition, showing WT or mutant R273H p53β-regulated DEGs in **(F)** U87-MG and A172 or **(G)** SF268 and SNB19 GBM cells. The indicated gene sets (p53β WT-regulated DEGs or p53β R273H-regulated DEGs) refer to genes regulated in the two WT or two R273H *TP53* GBM cell lines, respectively, upon expression of the corresponding p53β. **(H-K)** Gene set enrichment analysis (GSEA) plots showing enrichment scores (ES, upper panels) and ranked gene scores (lower panels) for **(H)** HALLMARK_P53_PATHWAY in U87-MG, **(I)** KEGG_ P53_SIGNALING_PATHWAY in A172, and **(J,K)** HALLMARK_*MYC*_TARGETS_V1 in SNB19 and SF268, respectively. Normalized enrichment scores (NES) and adjusted p values are indicated (adj p <0.05). **(L,M)** L2P pathway analysis of Hallmark gene sets comparing EV and p53β DOX vs DMSO contrasts in **(L)** WT or **(M)** mutant R273H *TP53* GBM cell lines. Dot color indicates enrichment score, dot size represents -$log_{10}$(FDR), and circles and crosses indicate FDR <0.05 and FDR>0.05, respectively. RNA-seq analyses were performed using three independent biological replicates per condition (n=3).

### Supplementary Figure 4

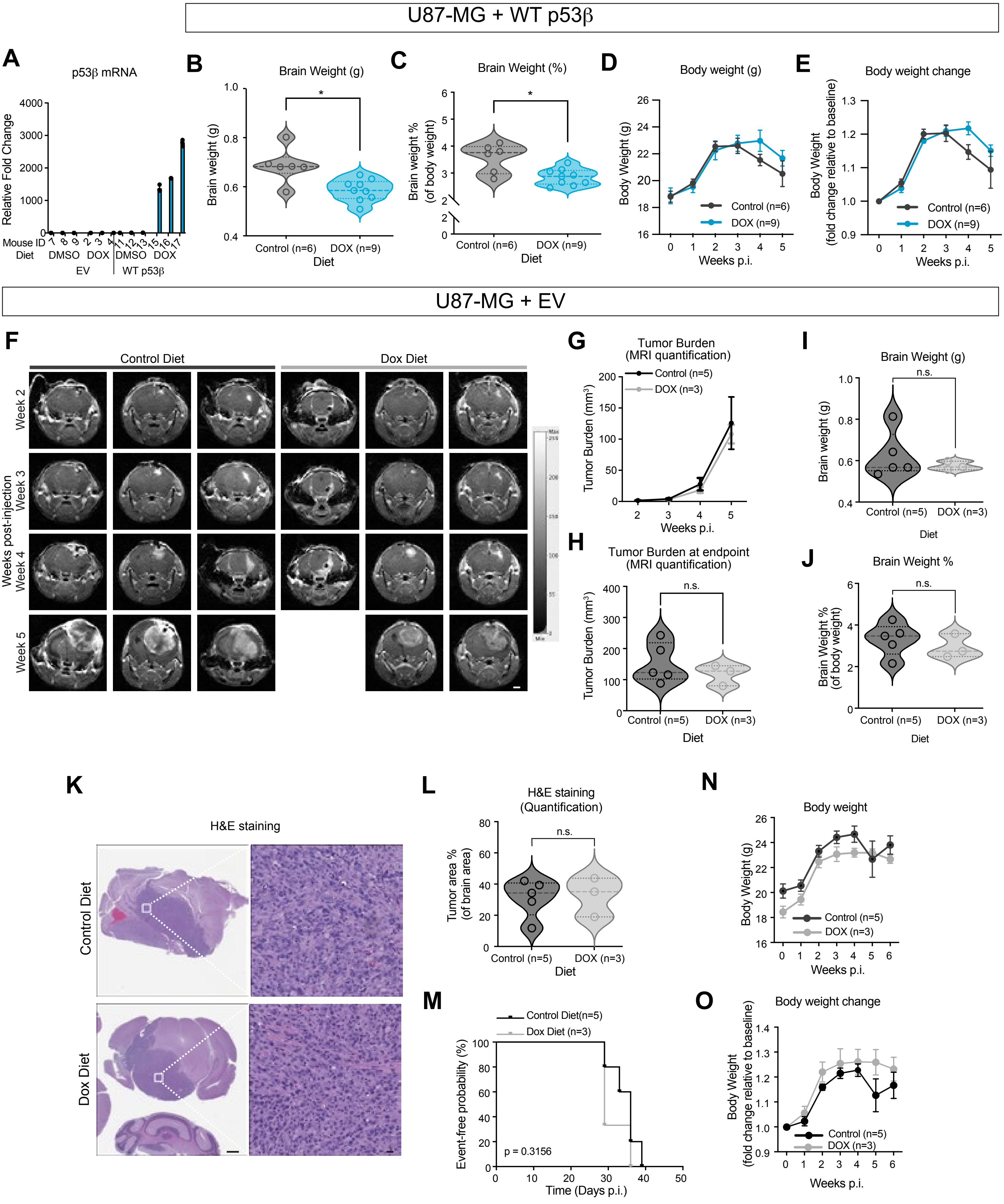

**<u>Supplementary Figure 4:</u> Doxycycline-induced WT p53β, but not an EV control, suppresses tumor growth in an intracranial U87-MG GBM xenograft model.** NSG mice were intracranially injected with human U87-MG cells stably expressing doxycycline-inducible WT p53β **(B-E,** control, n=6; DOX, n=9**)** or an EV control **(F-O**, control, n=5; Dox, n=3, unless otherwise specified). This experimental design is described in Figure 5A. **(A)** Relative p53β mRNA expression normalized to GAPDH in mice injected with U87-MG cells expressing EV control or WT p53β under DOX or control diet (n=3, per group). **(B, I)** Brain weight at endpoint (necropsy). Statistical significance was assessed by Welch's t-test: **(B)** p=0.0162; **(I)** p=0.3774. **(C, J)** Brain weight expressed as a percentage of body weight at necropsy. Statistical significance was assessed by Welch's t-test: **(C)** p=0.0194; **(J)** p=0.4681. **(D, N)** Longitudinal body weight (g) at the indicated times post-injection. **(E, O)** Relative body weight (fold change from baseline) at the indicated times post-injection. **(F)** Representative MRI images of mice bearing EV control tumors obtained at weeks 2 through 5 post-injection. Scale bar = 2 mm. **(G)** MRI-based quantification of tumor burden ($mm^3$) in EV control mice at the indicated times post-injection. Statistical significance was assessed by two-way ANOVA followed by Sidak's multiple comparisons test **(**week 4, p=0.9916; week 5, p=0.9298). **(H)** MRI-based quantification of tumor burden ($mm^3$) in EV control mice at endpoint. Statistical significance was assessed by Welch's t-test (p=0.8877). **(K)** Representative hematoxylin and eosin (H&E)-stained brain sections (left) and higher magnification images of the boxed regions (right). Scale bars: 1 mm (left); 20 μm (right). **(L)** Quantification of tumor area as a percentage of total brain area in H&E-stained sections. Statistical significance was assessed by Welch's t-test, p=0.8905. **(M)** Kaplan-Meier curve showing the event-free probability in mice bearing EV control tumors, with events defined as in Figure 5G. Statistical significance was assessed by the log-rank (Mantel-Cox) test (p=0.5284). Individual data points represent individual mice (B, C, H-J, L). *p < 0.05; **p<0.01; ***p<0.001; ****p<0.0001. ns, not significant.

### Supplementary Figure 5

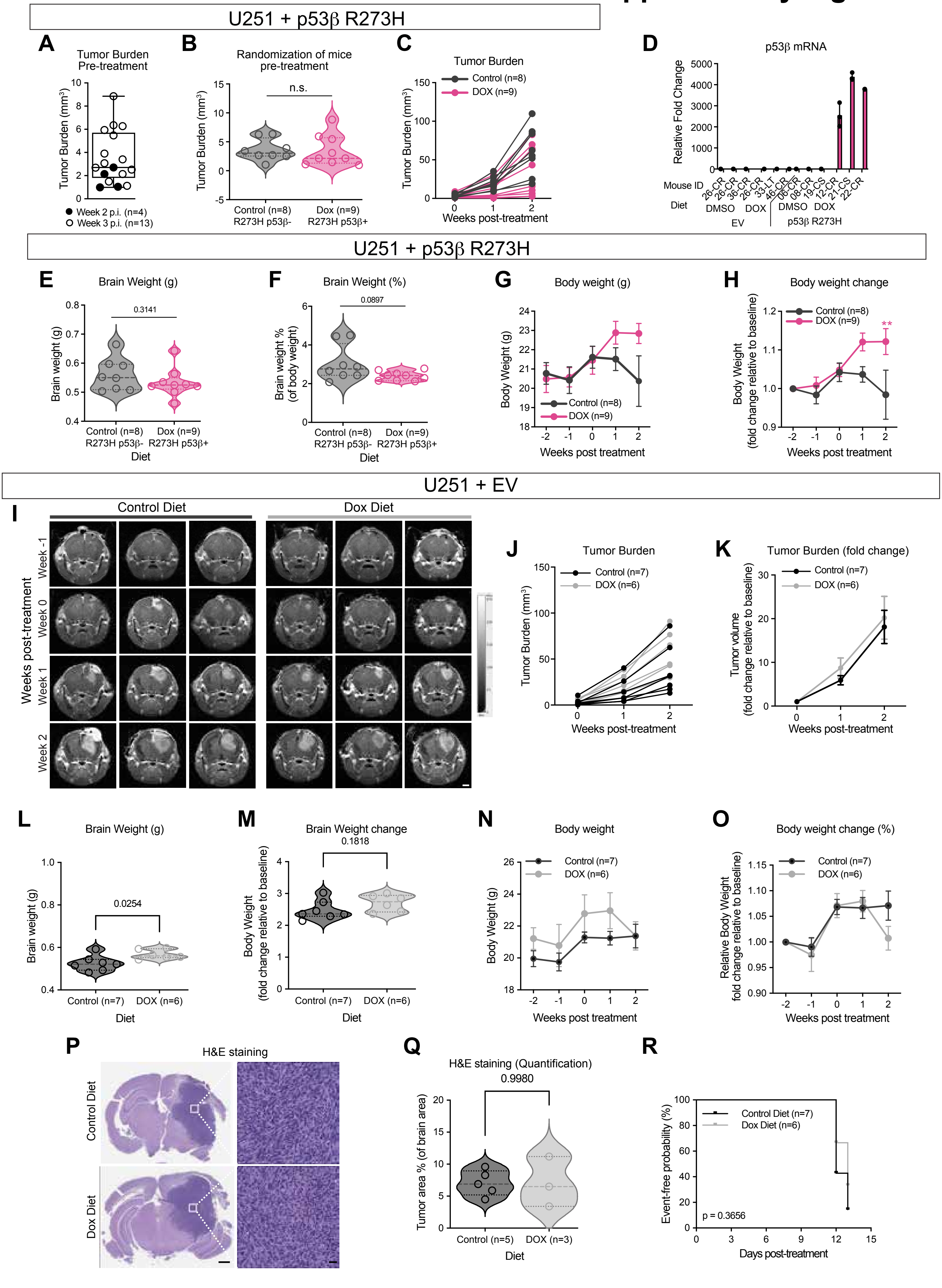

__Supplementary Figure 5:__ **Mutant p53β R273H suppresses intracranial GBM tumor growth, whereas doxycycline has no effect in EV control tumors.** NSG mice were intracranially injected with human U251 GBM cells stably expressing doxycycline-inducible mutant p53β R273H (**A-C, E-H**; control, n=8; DOX, n=9, unless indicated**)** or an EV control **(I-R**; control, n=7; DOX, n=6, unless indicated). The experimental design is described in Figure 6A. **(A)** Pretreatment tumor burden ($mm^3$) measured by MRI at weeks 2 (n=4) or 3 (n=13) post-injection. **(B)** Baseline tumor burden ($mm^3$) at week 0, showing the distribution of mice randomized to control (n=8) or DOX (n=9) diets. Statistical significance was assessed by Mann-Whitney t-test (p=0.4807). **(C)** MRI-based quantification of tumor burden ($mm^3$) in individual mice at the indicated times post-treatment. (Control, n=8; DOX, n=9). **(D)** Relative p53β mRNA expression normalized to GAPDH in mice injected with U251 cells expressing EV control or mutant R273H p53β under DOX or control diet (n=3, per group). **(E, L)** Brain weight at endpoint (necropsy) in mice bearing **(E)** mutant p53β R273H or **(L)** EV control tumors. (Welch's t-test: (E) p=0.3141; (L) p=0.0254). **(F, M)** Brain weight expressed as a percentage of body weight at necropsy (Welch's t-test: (F) p=0.0897; (M) p=0.1818) **(G, N)** Longitudinal body weight (g) at the indicated times post-treatment in mice bearing **(G)** mutant p53β R273H or **(N)** EV control tumors. **(H, O)** Relative body weight (fold change from week 0) at the indicated times post-treatment in mice bearing **(H)** mutant p53β R273H or **(O)** EV control tumors. **(I)** Representative MRI images of mice bearing EV control tumors obtained at the indicated times post-treatment. Scale bar = 2 mm. **(J)** MRI-based quantification of tumor volume ($mm^3$) for individual mice at the indicated times post-treatment. **(K)** MRI-based quantification of tumor burden expressed as fold change relative to each animal's pretreatment tumor volume (baseline, week 0). Statistical significance was assessed by two-way ANOVA followed by Sidak's multiple comparisons test (Week 0, p>0.9999; week 1, p=0.8527; week 2, p=0.9279). **(P)** Representative hematoxylin and eosin (H&E)-stained brain sections (left) and higher magnification images of the boxed regions (right). Scale bars: 1 mm (left), 20 µm (right). **(Q)** Quantification of tumor area as a percentage of total brain area in H&E-stained sections. Only mice with pretreatment tumor volumes of < 3.5 $mm^3$ (average across all EV and mutant R273H p53β groups) were included. (Control, n=5; DOX, n=3). Welch's t-test, p=0.9980. **(R)** Kaplan-Meier curve showing event-free probability, with an event defined as the first occurrence of any of the following: 10% body weight loss, tumor growth rate>19 (average across groups), or onset of neurological signs (hunched posture, head tilt, circling). Log-rank (Mantel-Cox) test, p=0.3656. Longitudinal data (G, H, K, N, O) are presented as mean +/- SEM. Individual data points represent individual mice (A-F, J, L, M, Q). ns, not significant.

### Supplementary Figure 6

#### Full Size Western Blots

##### Supplementary Figure 1D

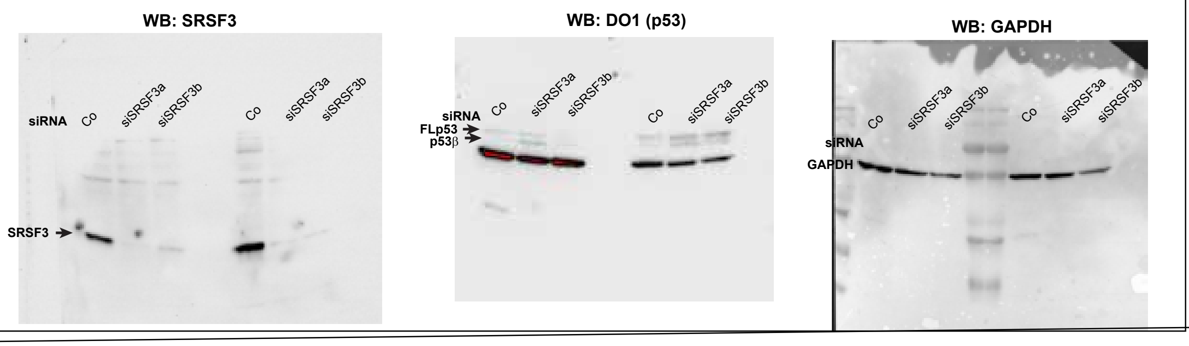


##### Figure 1B

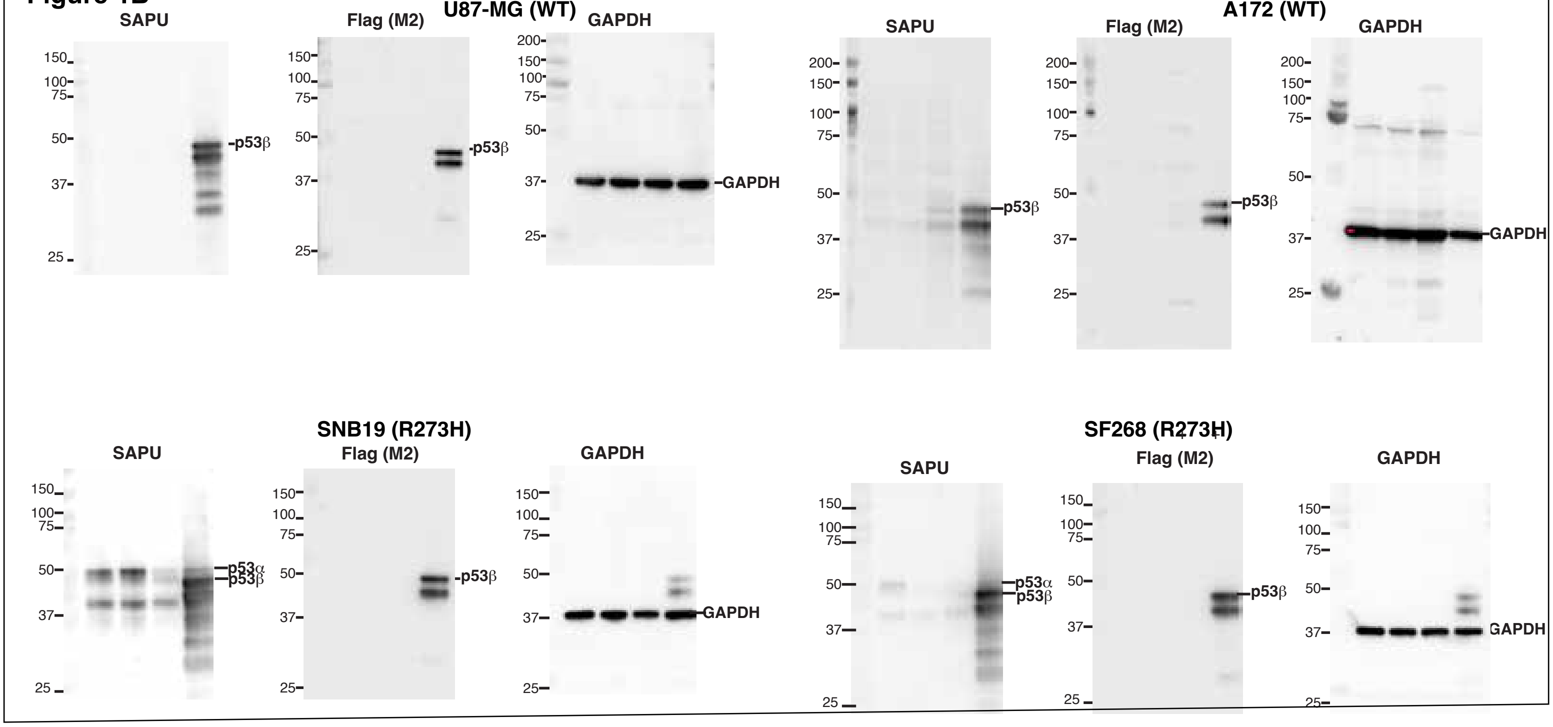


##### Figure 2E

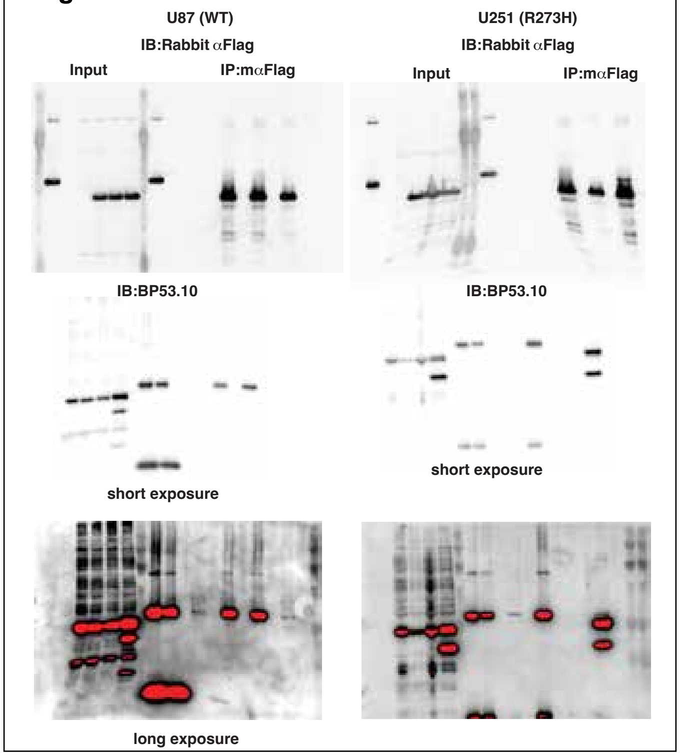


##### Figure 4D

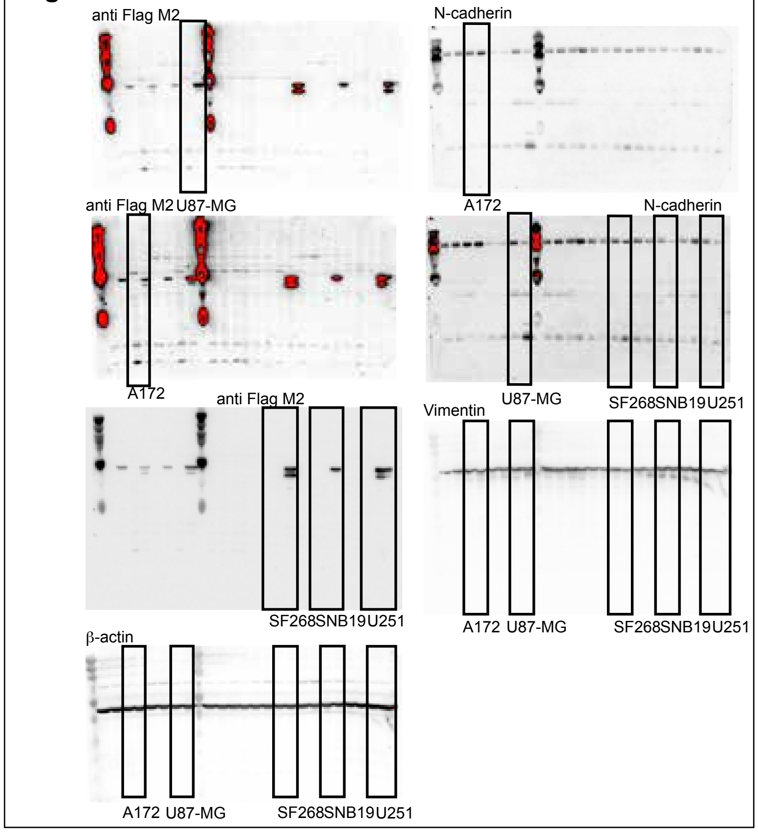

**Supplementary Figure 6:** Full, uncropped western blots corresponding to the indicated supplementary or main figures.

#### Supplementary Table S1. Key resources used in this study

##### A. Cell lines, siRNAs, and expression constructs

| Category | Resource | Manufacturer/source | Identifier/Catalog # | Application/notes |
|---|---|---|---|---|
| Cell line | U87-MG | ATCC | CD Genomics (2022) | Human GBM; *TP53* WT |
| Cell line | A172 | ATCC | CD Genomics (2022) | Human GBM; *TP53* WT |
| Cell line | SF268 | NCI DCTD repository | CD Genomics (2022) | Human GBM; *TP53* R273H |
| Cell line | SNB19 | NCI DCTD repository | CD Genomics (2022) | Human GBM; *TP53* R273H |
| Cell line | U251 | NCI DCTD repository | CD Genomics (2022) | Human GBM; *TP53* R273H |
| Cell line | LNZ308N | ATCC | CD Genomics (2022) | Human GBM; *TP53* -null |
| siRNA | SRSF3-targeting siRNA 1 | Thermo Fisher Scientific | s12732 | SRSF3 knockdown |
| siRNA | SRSF3-targeting siRNA 2 | [TO COMPLETE] | s12733 | Second independent siRNA |
| siRNA | Negative control siRNA | Thermo Fisher Scientific | 4390843 | Nonspecific control |
| Lentiviral vector | pLV_Tet3G | VectorBuilder | VB230221-1497mpj | Tet-On transactivator |
| Lentiviral vector | pLV_TRE3G_Empty_Vector | VectorBuilder | VB230313-1204zyv | Empty-vector control |
| Lentiviral vector | pLV_TRE3G_Flag_p53β_WT | VectorBuilder | VB230314-1334atb | DOX-inducible WT p53β |
| Lentiviral vector | pLV_TRE3G_Flag_p53β_R273H | VectorBuilder | VB230314-1331zfw | DOX-inducible mutant p53β R273H |
| Expression plasmid | WT p53α | J-C Bourdon's lab | {Marcel, 2014 #22} | Reconstitution, luciferase and co-IP |
| Expression plasmid | Mutant p53α R273H | J-C Bourdon's lab | {Marcel, 2014 #22} | Reconstitution, luciferase and co-IP |
| Expression plasmid | pcDNA control | J-C Bourdon's lab | {Marcel, 2014 #22} | Negative control |
| Reporter plasmid | p21 promoter–Firefly luciferase | J-C Bourdon's lab | {Marcel, 2014 #22} | p21 promoter activity |
| Reporter plasmid | Renilla luciferase control | J-C Bourdon's lab | {Marcel, 2014 #22} | Reporter normalization |

##### B. Antibodies and immunoprecipitation reagents

| Category | Resource | Manufacturer/source | Catalog number/reference | Application |
|---|---|---|---|---|
| Antibody | DO-1 (p53 isoforms) | Santa Cruz Biotechnology | sc-126 | Western blot |
| Antibody | SAPU (pantromic, p53 isoforms) | Published reagent | {Khoury, 2010 #50} | Western blot |
| Antibody | BP.53 (p53alfa-specific) | Santa Cruz Biotechnology | sc-263 | Western blot |
| Antibody | SRSF3 | Santa Cruz Biotechnology | sc-13510 | Western blot |
| Antibody | GAPDH | EMD Millipore | ABS16 | Western blot |
| Antibody | Anti-FLAG M2 | Sigma-Aldrich | F3125 | Western blot |
| Antibody | N-cadherin | Cell Signaling Technology | 4061 | Western blot |
| Antibody | Vimentin | Abcam | ab137321 | Western blot |
| Antibody | β-actin | Santa Cruz Biotechnology | 47778 | Western blot |
| Antibody | p21 | Cell Signaling Technology | 2947S | Immunofluorescence |
| Antibody | Ki67 | Abcam | ab15580 | Immunofluorescence |
| Antibody | Cleaved caspase-3 | Cell Signaling Technology | 9661S | Immunofluorescence |
| IP reagent | Anti-FLAG M2 Affinity Gel | Sigma-Aldrich | A2220 | Co-immunoprecipitation |
| IP reagent | 3×FLAG peptide | Sigma-Aldrich | F4799 | Elution of FLAG-tagged proteins |

#### C. Quantitative RT-PCR assays and primers

| Target | Detection method | Manufacturer | Assay ID or forward primer (5'–3') | Reverse primer (5'–3') |
|---|---|---|---|---|
| *CDKN1A* (p21) | TaqMan | Thermo Fisher Scientific | Hs00355782_m1 | — |
| *BAX* | TaqMan | Thermo Fisher Scientific | Hs00180269_m1 | — |
| *BBC3* (PUMA) | TaqMan | Thermo Fisher Scientific | Hs00248075_m1 | — |
| GAPDH | TaqMan | Thermo Fisher Scientific | Hs02758991_g1 | — |
| *SRSF3* | SYBR Green | Invitrogen primers | AGCTGATGCAGTCCGAGAG | GGTGGGCCACGATTTCTA |
| *p53β* | SYBR Green | Invitrogen primers | GCGAGCACTGCCCAACA | GAAAGCTGGTCTGGTCCTGAA |

| | | | | |
|---|---|---|---|---|
| *GAPDH* | SYBR Green | Invitrogen primers | CCATCTTCCAGGAGCGAGA | TGTCATACCAGGAAATGAGC |
| *Total TP53 (exon 8)* | SYBR Green | Invitrogen primers | CCTCAGCATCTTATCCGAGTGG | TGGATGGTGGTACAGTCAGAGC |

#### D. Commercial kits, reagents, and culture materials

| **Category** | **Resource** | **Manufacturer/source** | **Catalog number** | **Application** |
|---|---|---|---|---|
| RNA extraction | RNeasy Mini Kit | QIAGEN | 74104 | qRT-PCR |
| RNA extraction | RNeasy Plus Kit | QIAGEN | 74134 | RNA-seq |
| Reverse transcription | High-Capacity cDNA Reverse Transcription Kit | Thermo Fisher Scientific | 4368813 | cDNA synthesis |
| qPCR | Taqman | Thermo Fisher Scientific | 4304437 | Quantitative PCR |
| Senescence assay | SA-β-galactosidase Staining Kit | Cell Signaling Technology | 9860 | Senescence detection |
| Transfection | Lipofectamine RNAiMAX | Thermo Fisher Scientific | 13778150 | siRNA delivery |
| Transfection | TurboFect | Invitrogen | R0531 | Co-IP plasmid transfection |
| Reporter assay | Dual-Luciferase Reporter Assay System | Promega | E1910 | Firefly/Renilla detection |
| IL-6 assay | Human IL-6 ELISA Kit | Sigma-Aldrich | EZIL6 | Secreted IL-6 measurement |
| 3D culture | Tumorsphere MediumXF | Promocell | C-28070 | Spheroid experiments |
| Invasion reagent | Matrigel | Corning | 354234 | Spheroid invasion |
| Selection antibiotic | Puromycin | Sigma-Aldrich | P9620 | Lentiviral selection |
| Selection antibiotic | Blasticidin | Sigma-Aldrich | SBR00022 | Lentiviral selection |
| Induction reagent | Doxycycline hyclate | Sigma-Aldrich | D3072 | Tet-On expression |
| Mounting medium | VECTASHIELD with DAPI | Vector Laboratories | H-1200-10 | Immunofluorescence |
| Animal model | Female NSG mice, 6–8 weeks | Jackson laboratories | NOD.Cg-Prkdc^scid^ Il2rg^tm1Wjl^/SzJ | Intracranial xenografts |
| Animal diet | Doxycycline-supplemented diet | Bio-Serve | S3888 | 200 mg/kg diet |

| | | | | |
|---|---|---|---|---|
| Animal diet | Doxycycline-supplemented diet | Bio-Serve | S4207 | |
| Animal diet | Standard chow | Purina | 5L79 | Routine feeding |

**Supplementary Table S2. Animal allocation and analysis cohorts for orthotopic GBM xenograft studies**

| Mouse model | Inducible construct | # injected | Control diet (n) | DOX diet (n) | Dox/control diet randomization | Exclusions |
|---|---|---|---|---|---|---|
| U87-MG (*TP53* WT) | WT p53β | 15 | 6 | 9 | Week 1 post-injection | None |
| U87-MG (*TP53* WT) | Empty vector (EV) | 10 | 5 | 3 | Week 1 post-injection | 2 animals died post surgery or during MRI |
| U251 (*TP53* R273H) | Mutant p53β R273H | 12 | 4 | 4 | After MRI-confirmed engraftment at week 2 or 3 | 4 animals were excluded due to death post-surgery, no tumor growth at baseline, or tumor engraftment at a different area than expected |
| U251 (*TP53* R273H) | Mutant p53β R273H | 12 | 4 | 5 | After MRI-confirmed engraftment at week 2 or 3 | 3 animals were excluded due to death post-surgery, no tumor growth at baseline, or tumor engraftment at a different area than expected |
| U251 (*TP53* R273H) | Empty vector (EV) | 12 | 4 | 3 | After MRI-confirmed engraftment at week 2 or 3 | 5 animals were excluded due to death post-surgery, no tumor growth at baseline, or tumor engraftment at a different area than expected |
| U251 (*TP53* R273H) | Empty vector (EV) | 12 | 3 | 3 | After MRI-confirmed engraftment at week 2 or 3 | 6 animals were excluded due to death post-surgery, no tumor growth at baseline, or tumor engraftment at a different area than expected |
